# Modification of forests by people means only 40% of remaining forests have high ecosystem integrity

**DOI:** 10.1101/2020.03.05.978858

**Authors:** H.S. Grantham, A. Duncan, T. D. Evans, K. Jones, H. Beyer, R. Schuster, J. Walston, J. Ray, J. Robinson, M. Callow, T. Clements, H.M. Costa, A. DeGemmis, P.R. Elsen, J. Ervin, P. Franco, E. Goldman, S. Goetz, A. Hansen, E. Hofsvang, P. Jantz, S. Jupiter, A. Kang, P. Langhammer, W.F. Laurance, S. Lieberman, M. Linkie, Y. Malhi, S. Maxwell, M. Mendez, R. Mittermeier, N. Murray, H. Possingham, J. Radachowsky, C. Samper, J. Silverman, A. Shapiro, B. Strassburg, T. Stevens, E. Stokes, R. Taylor, T. Tear, R. Tizard, O. Venter, P. Visconti, S. Wang, J.E.M. Watson

**Affiliations:** Wildlife Conservation Society, Global Conservation Program, Bronx, New York, 10460 USA; School of Biological Sciences, University of Queensland, St. Lucia, Queensland, Australia; Department of Biology, 1125 Colonel By Drive, Carleton University, Ottawa ON, K1S 5B6 Canada; United Nations Development Programme, One United Nations Plaza, New York, NY, 10017, USA; World Resources Institute, Washington, DC, USA; Global Earth Observation & Dynamics of Ecosystems Lab, School of Informatics, Computing, and Cyber Systems, Northern Arizona University, Flagstaff, AZ, 86011, USA; Landscape Biodiversity Lab, Ecology Department, Montana State University, Bozeman, MT, 59717, USA; Rainforest Foundation Norway, Mariboes gate 8, 0183 Oslo; Global Wildlife Conservation, P.O. Box 129, Austin, Texas 78767, USA; School of Life Sciences, Arizona State University, P.O. Box 874501, Tempe, Arizona 85287, USA; Centre for Tropical Environmental and Sustainability Science, College of Science and Engineering, James Cook University, Cairns, QLD 4878, Australia; Environmental Change Institute, School of Geography and the Environment, University of Oxford, Oxford, United Kingdom; School of Earth and Environmental Sciences, University of Queensland, Brisbane, Australia; The Nature Conservancy, Arlington, VA, USA; World Wide Fund for Nature Germany, Space+Science; Natural Resource and Environmental Studies Institute, University of Northern British Columbia, Prince George, Canada; International Institute of Sustainability, Rio de Janeiro, 22460-320, Brazil; International Institute for Applied Systems Analysis, Laxenburg, Austria

## Abstract

Many global environmental agendas, including halting biodiversity loss, reversing land degradation, and limiting climate change, depend upon retaining forests with high ecological integrity, yet the scale and degree of forest modification remains poorly quantified and mapped. By integrating data on observed and inferred human pressures and an index of lost connectivity, we generate the first globally-consistent, continuous index of forest condition as determined by degree of anthropogenic modification. Globally, only 17.4 million km^2^ of forest (40.5%) have high landscape level integrity (mostly found in Canada, Russia, the Amazon, Central Africa and New Guinea) and only 27% of this area is found in nationally-designated protected areas. Of the forest in protected areas, only 56% has high landscape level integrity. Ambitious policies that prioritize the retention of forest integrity, especially in the most intact areas, are now urgently needed alongside current efforts aimed at halting deforestation and restoring the integrity of forests globally.

## Introduction

Deforestation is a major environmental issue ^1^, but far less attention has been given to the degree of anthropogenic modification of remaining forests, which reduces ecosystem integrity and diminishes many of the benefits that these forests provide ^2,3^. This is worrying since modification is potentially as significant as outright forest loss in determining overall environmental outcomes^4^. There is increasing recognition of this issue, for forests and other ecosystems, in synthesis reports by global science bodies e.g. ^5^, and it is now essential that the scientific community develop improved tools and data to facilitate the consideration of levels of integrity in decision-making. Mapping and monitoring this globally will provide essential information for coordinated global, national and local policy-making, planning and action, to help nations and other stakeholders achieve the Sustainable Development Goals (SDGs) and implement other shared commitments such as the United Nations Convention on Biological Diversity (CBD), Convention to Combat Desertification (UNCCD), and Framework Convention on Climate Change (UNFCCC).

Ecosystem integrity is foundational to all three of the Rio Conventions (UNFCCC, UNCCD, CBD). As defined by Parrish *et al.* ^6^, it is essentially the degree to which a system is free from anthropogenic modification of its structure, composition and function. Such modification causes the reduction of many ecosystem benefits, and is often also a precursor to outright deforestation ^7,8^. Forests largely free of significant modification (i.e. forests having high ecosystem integrity), typically provide higher levels of many forest benefits than modified forests of the same type ^9^, including; carbon sequestration and storage ^10^, healthy watersheds ^11^, traditional homelands for imperiled cultures ^12^, contribution to local and regional climate processes ^13^, and forest-dependent biodiversity ^14-17^. Industrial-scale logging, fragmentation by infrastructure, farming (including cropping and ranching) and urbanization, as well as less visible forms of modification such as over-hunting, wood fuel extraction and changed fire or hydrological regimes ^18,19^, all degrade the degree to which forests still support these benefits, as well as their long-term resilience to climate change ^9^. There can be trade-offs however, between the benefits best provided by less-modified forests (e.g., regulatory functions such as carbon sequestration) and those production services that require some modification (e.g., timber production). These trade-offs can, at times, result in disagreement among stakeholders as to which forest benefits should be prioritized ^20^.

In recent years, easily accessible satellite imagery and new analytical approaches have dramatically improved our ability to map and monitor forest extent globally ^21-23^. However, while progress has been made in developing tools for assessment of global forest losses and gains, consistent monitoring of the degree of forest modification has proved elusive ^24,25^. Technical challenges include the detection of low intensity and unevenly distributed forest modification, the wide diversity of changes that comprise forest modification, and the fact that many changes are concealed by the forest canopy ^24^. New approaches are emerging on relevant forest indicators, such as canopy height, canopy cover and fragmentation, and maps of different human pressures, which are used as proxies for impacts on forests e.g., ^26,27,28^. Some binary measures of forest modification, such as Intact Forest Landscapes ^29^ and wilderness areas ^30^, have also been mapped at the global scale and used to inform policy, but do not resolve the degree of modification within remaining forests, which we aimed to do with this assessment.

Human activities influence the integrity of forests at multiple spatial scales, including intense, localized modifications such as road-building and canopy loss, more diffuse forms of change that are often spatially associated with these localized pressures (e.g. increased accessibility for hunting, other exploitation, and selective logging), and changes in spatial configuration that alter landscape-level connectivity. Previous studies have quantified several of these aspects individually e.g. ^26,27,28^, but there is a need to integrate them to measure and map the overall degree of modification considering these landscape-level anthropogenic influences at each site. Here, we integrate data on forest extent defined as all woody vegetation taller than 5 m, following ^22^, ‘observed’ human pressures (e.g. infrastructure) which can be directly mapped using current datasets, other ‘inferred’ human pressures (e.g. collection of forest materials) that occur in association with those that are observed but cannot be mapped directly, and alterations in forest connectivity, to create the “Forest Landscape Integrity Index” (FLII), that describes the degree of forest modification for the beginning of 2019 (Fig. 1). The result is the first globally applicable, continuous-measure map of landscape level forest integrity (hereafter, integrity), which offers a timely indicator of the status and management needs of Earth’s remaining forests, as well as a flexible methodological framework (Fig. 1) for measuring changes in forest integrity that can be adapted for more detailed analysis at national or subnational scales.

**Figure 1.**
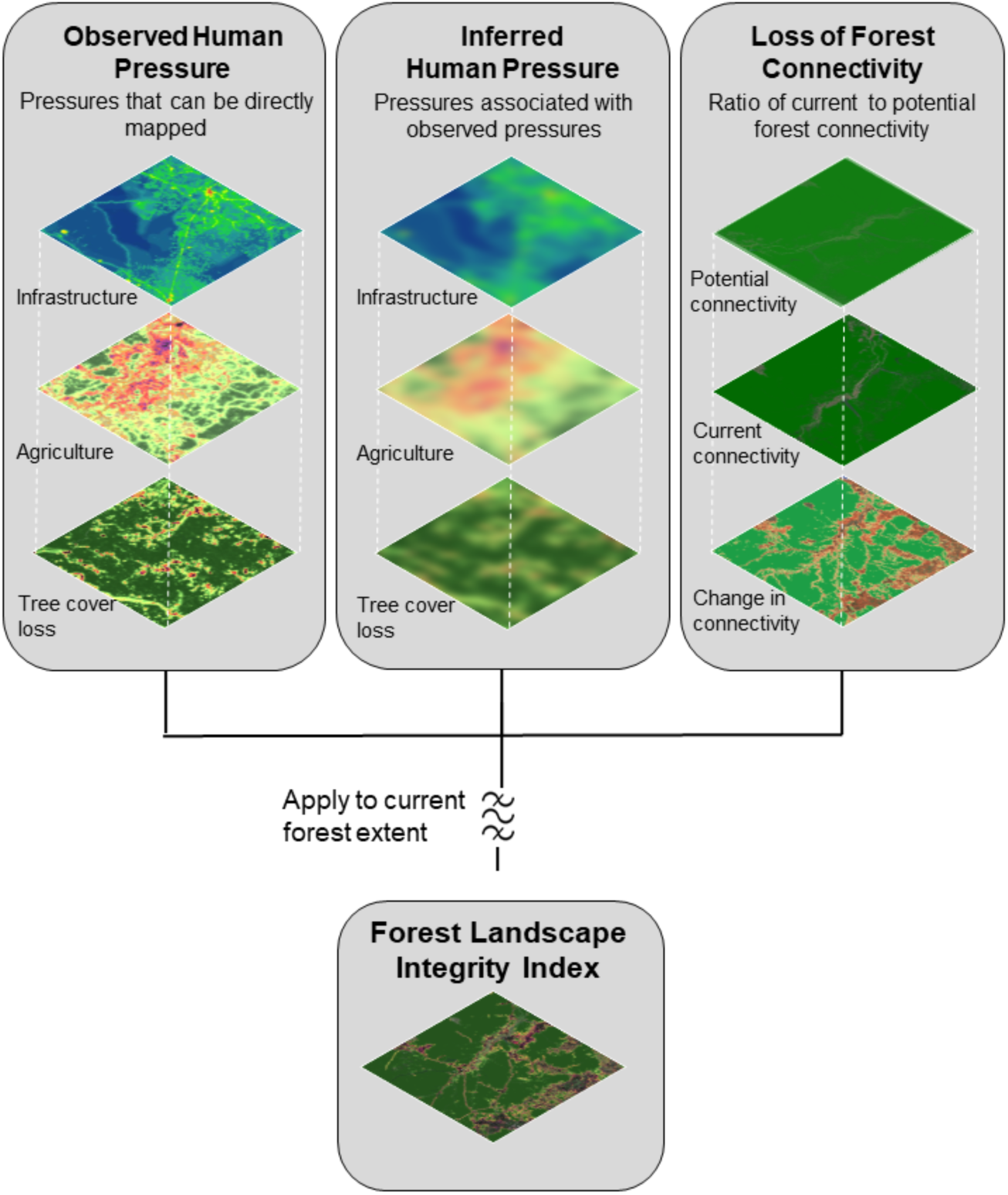
The Forest Landscape Integrity Index was constructed based on three main data inputs: 1) observed pressures (infrastructure, agriculture, tree cover loss), 2) inferred pressure modelled based on proximity to the observed pressures, and 3) change in forest connectivity.

## Results

Forest modification caused by human activity is both highly pervasive and highly variable across the globe (Fig. 2). We found 31.2% of forests worldwide are experiencing some form of ‘observed’ human pressure. Our models also inferred the likely occurrence of other pressures, and the impacts of lost connectivity, in almost every forest location (91.2% of forests), albeit sometimes at very low levels. Diverse, recognizable patterns of forest integrity can be observed in our maps at a range of scales, depending on the principal forms and general intensity of human activity in an area. Broad regional trends can be readily observed, for example the overall gradient of decreasing human impact moving northwards through eastern North America (Fig. 2), and finer patterns of impact are also clearly evident, down to the scale of individual protected areas, forest concessions, settlements and roads (Fig. S2).

**Figure 2.**
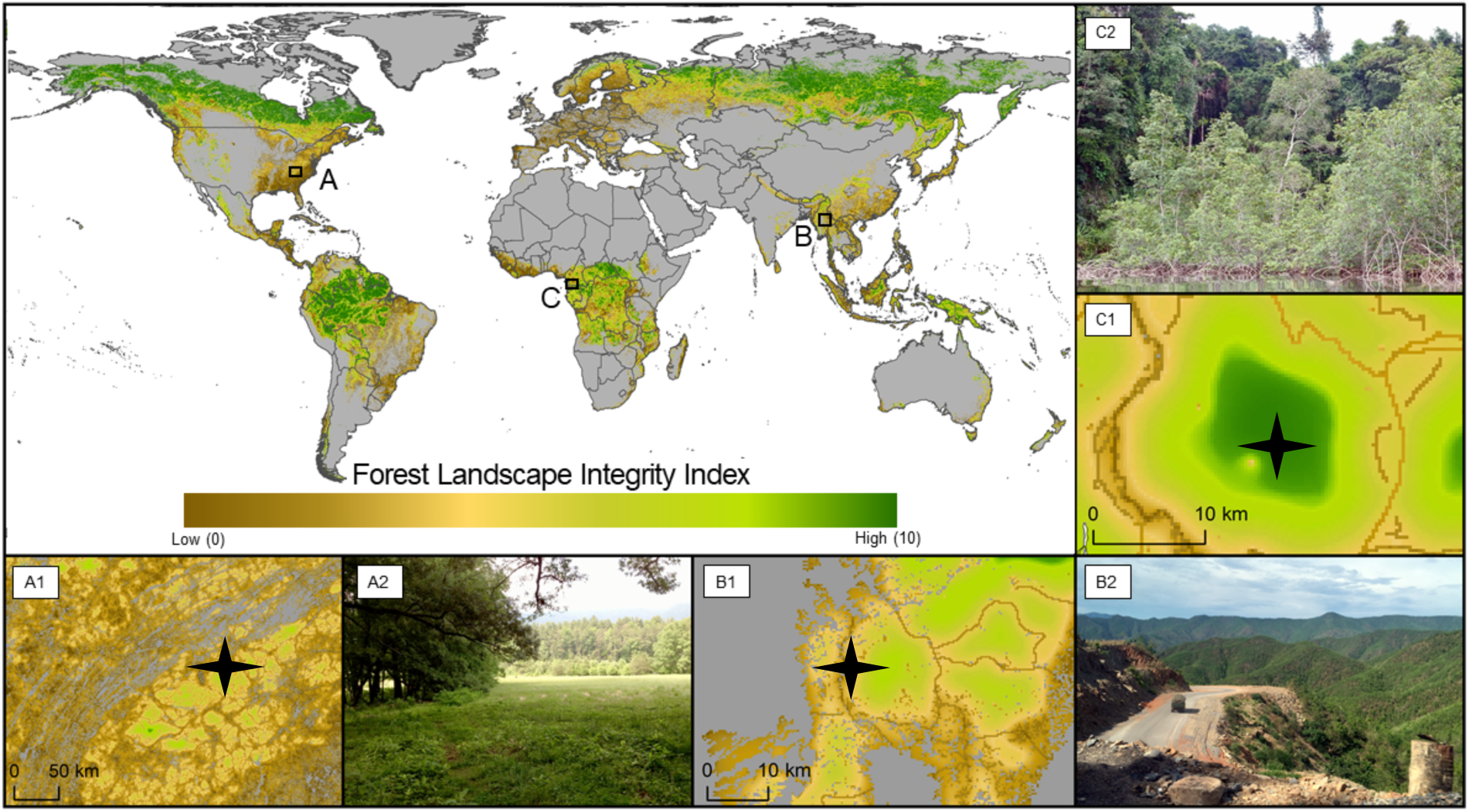
A global map of Forest Landscape Integrity for 2019. Three regions are highlighted including A) USA, B) Equatorial Guinea C) Myanmar. For a) shows the edge of Smoky Mountains National Park in Tennessee b) shows a logging truck passing through some partially degraded forest along a newly constructed highway in Shan State, c) shows an intact mangrove forest within Reserva Natural del Estuario del Muni, near the border with Gabon. The stars indicate approximately where the photos were taken (A2, B2 and C2).

FLII scores range from 0 (lowest integrity) to 10 (highest). We discretized this range to define three broad illustrative categories: low (≤6.0); medium (>6.0 and <9.6); and high integrity (≥9.6) by benchmarking against reference locations worldwide (see Methods). Only 40.5% (17.4 million km^2^) of forest was classified as having high integrity (Fig. 3; Table 1). Moreover, even in this category of high integrity (36%) still showed at least a small degree of human modification. The remaining 59% (25.6 million km^2^) of forest was classified as having low or medium integrity, including 25.6% (11 million km^2^) with low integrity (Fig. 3; Table 1). When we analyzed across biogeographical realms defined by ^31^ not a single biogeographical realm of the world had more than half of its forests in the high category (Fig. 3; Table 1).

**Table 1.** A summary of the Forest Landscape Integrity Index scores for each biogeographic realm globally, measuring the mean score, in addition to the area and proportion of realm for each category of integrity. Scores are divided into three categories of integrity: high, medium and low.

| Biogeographic Realm | Total forest | FLII | High (9.6 - 10) |  | Medium (6 – 9.6) |  | Low (0 - 6) |  |
| --- | --- | --- | --- | --- | --- | --- | --- | --- |
|  | *Km<sup>2</sup>* | *Mean* | *Km<sup>2</sup>* | *% of realm* | *Km<sup>2</sup>* | *% of realm* | *Km<sup>2</sup>* | *% of realm* |
| Afrotropic | 7,362,740 | 7.34 | 2,450,953 | 33.3 | 2,903,483 | 39.4 | 2,008,304 | 27.3 |
| Australasia | 1,711,684 | 8.05 | 656,701 | 38.4 | 753,188 | 44.0 | 301,796 | 17.6 |
| Indo-malayan | 3,596,249 | 5.9 | 420,977 | 11.7 | 1,599,049 | 44.5 | 1,576,223 | 43.8 |
| Neotropic | 10,271,519 | 7.81 | 4,579,406 | 44.6 | 3,122,706 | 30.4 | 2,569,407 | 25.0 |
| Oceania | 23,389 | 7.66 | 5,279 | 22.6 | 14,331 | 61.3 | 3,780 | 16.2 |
| Palearctic | 12,172,668 | 8 | 5,571,997 | 45.8 | 3,910,629 | 32.1 | 2,690,042 | 22.1 |
| Nearctic | 7,794,117 | 7.84 | 3,716,855 | 47.7 | 2,257,518 | 29.0 | 1,819,744 | 23.3 |
| **Total** | 42,932,367 | 7.76 | 17,402,170 |  | 14,560,903 |  | 10,969,294 |  |

**Figure 3.**
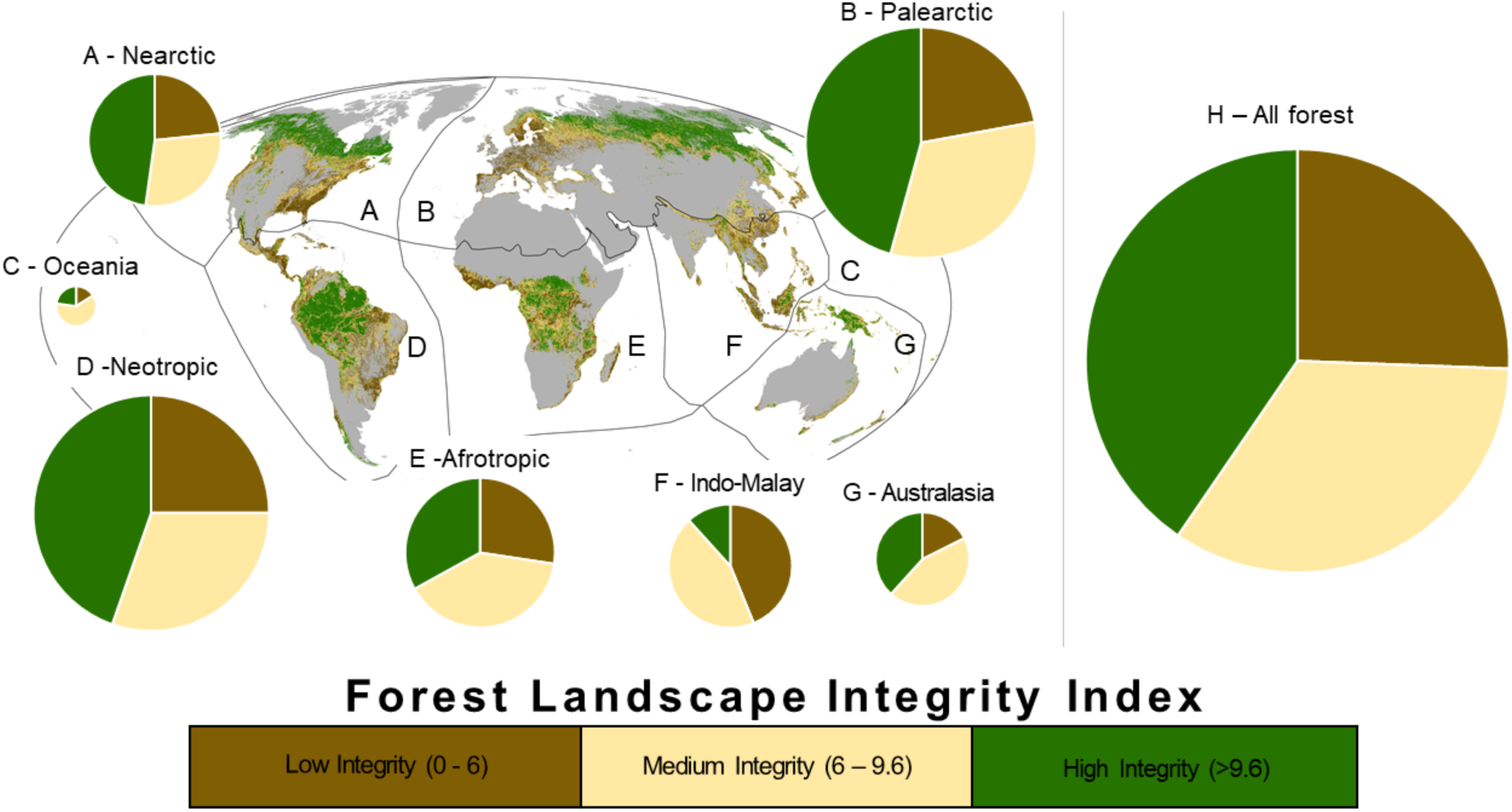
The Forest Landscape Integrity Index for 2019 categorized into three broad, illustrative classes and mapped for across each biogeographic realm (A – G). The size of the pie charts indicates the relative size of the forests within each realm (A - G), and H shows all the world’s forest combined.

The biogeographical realms with the largest area of forest with high integrity are the Paleartic, particularly northern Russia, and the Neartic, in northern Canada, and Alaska. There are also large areas of forest with high integrity in the Neotropics, concentrated in the Amazon region, including within the Guianas (Fig. 3, Table 1). The Afrotropic realm has significant areas with high integrity, particularly within the humid forests of central Africa (e.g., in Republic of Congo and Gabon) and in some of the surrounding drier forest/woodland belts (e.g., in South Sudan, Angola and Mozambique) (Fig. 3). In tropical Asia, the largest tracts of forest with high integrity are in New Guinea. Smaller but still very significant tracts of forest with high integrity are also scattered elsewhere in each of the main forested regions, including parts of Sumatra, Borneo, Myanmar and other parts of the greater Mekong subregion, Madagascar, West Africa, Mesoamerica, the Atlantic forests of Brazil, southern Chile, the Rocky Mountains, northern Assam, the Pacific forests of Colombia, the Caucasus, and the Russian Far East (Fig. 3).

Concentrations of forest with low integrity are found in many regions including west and central Europe, the south-eastern USA, island and mainland South-East Asia west of New Guinea, the Andes, much of China and India, the Albertine Rift, West Africa, Mesoamerica and the Atlantic Forests of Brazil (Fig. 3). The overall extent of forests with low integrity is greatest in the Paleartic realm, followed by the Neotropics, which are also those biogeographic realms with the largest forest cover (Table 1). The Indo-Malayan realm has the highest percentage with low integrity, followed by the Afrotropics (Fig. 3; Table 1).

These patterns result in variation with forest integrity scores in ways that allow objective comparisons to be made between locations and at a resolution relevant for policy and management planning, such as at national and sub-national scales. The global average FLII score for all countries is 5.48, representing generally low forest integrity, and a quarter of forested countries have a national average score < 4. National mean scores vary widely, ranging from >9 in Guyana, French Guiana, Gabon, Sudan and South Sudan to <3 in Sierra Leone and many west European countries (see Fig 4. and Table S5 for full list of countries). Provinces and other sub-national units vary even more widely (see Fig. S2 and Table S6)

**Figure 4.**
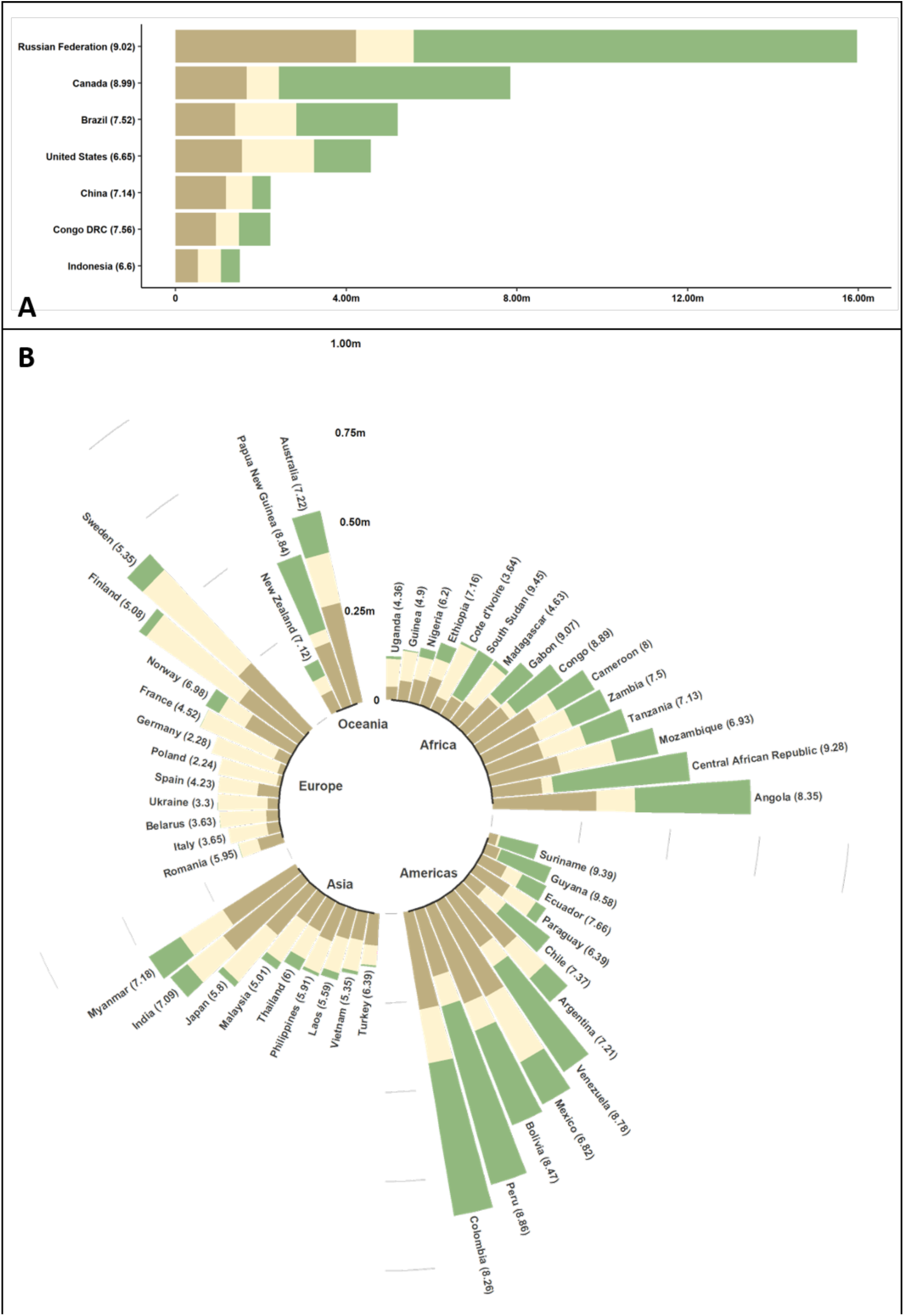
The Forest Landscape Integrity Index for 2019 categorized into three broad, illustrative classes for each major forested country in the world. (A) countries with a forest extent larger than 1 million km^2^, and (B) countries with forest extent between 1 million km^2^ and 100,000 km^2^ of forest. The size of the bar represents the area of a country’s forests.

Over one-quarter (26.1%) of all forests with high integrity fall within protected areas, compared to just 13.1% of low and 18.5% of medium integrity forests respectively. For all forests that are found within nationally designated protected areas (around 20% of all forests globally), we found the proportions of low, medium and high integrity forests were 16.8%, 30.3%, and 52.8% respectively (Table 2). Within the different protected area categories, we typically found that there was more area within the high integrity category versus the medium and low except for Category V (protected landscape/seascape) (Table 2). However, with 47.1% of forests within protected areas having low to medium integrity overall, it is clear that forests considered ‘protected’ are already often fairly modified (Table 2). Even though they are quite modified, some of these forests might still have high conservation importance, such as containing endangered species.

**Table 2.**
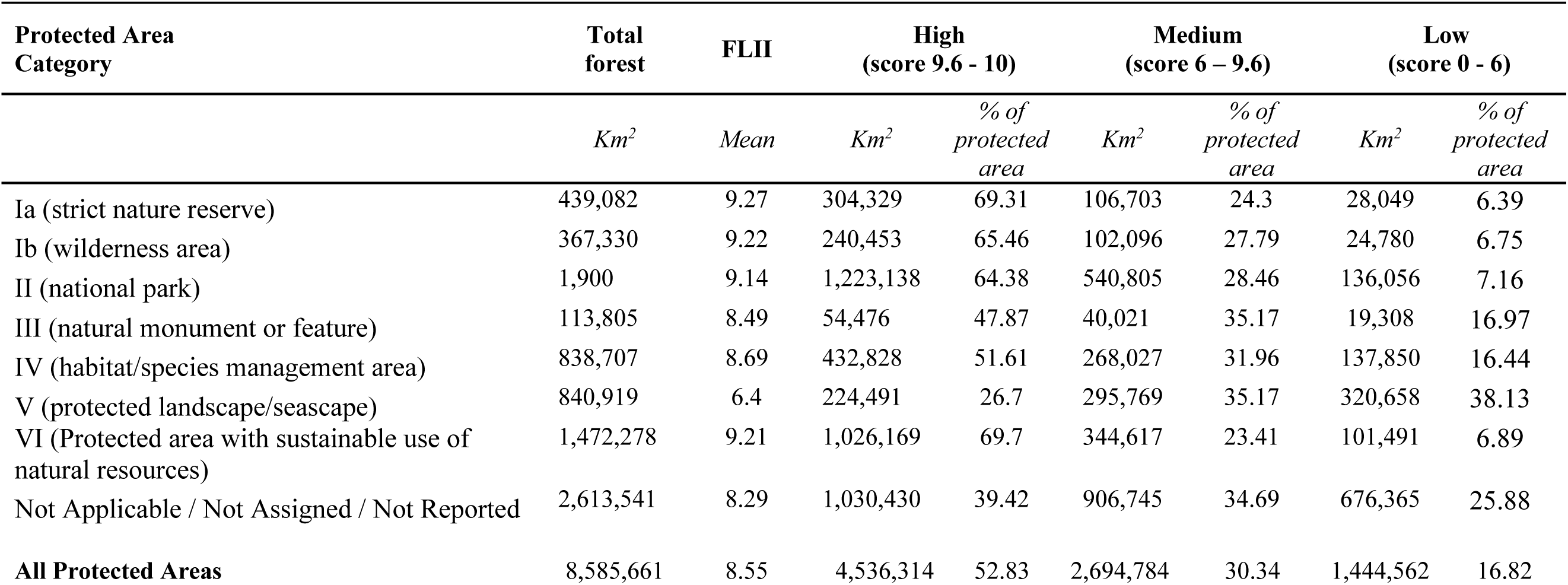
A summary of the Forest Landscape Integrity Index scores for each type of protected area designation based on the IUCN Protected Areas categories measuring mean score, in addition to the area and proportion of realm for each category of integrity. Scores are divided into three categories of integrity: high, medium and low.

## Discussion

By providing a transparent and defensible methodological framework, and by taking advantage of global data on forest extent, human drivers of forest modification, and changes in forest connectivity, our analysis paints a new, sobering picture of the extent of human impacts on the world’s forests. This analysis enables the changes that degrade many forest values *(8)* to be visualized in a new and compelling way and for policy makers and decision makers to see where forests that survive in good condition are found. By integrating data on multiple pressures that are known to modify forests, our analysis is the first to move global quantification beyond the use of simple categories, or solely using pressure indicators as proxies for integrity, to a more nuanced depiction of this issue as a continuum, recognising that not all existing forests are in the same condition. Our analysis reveals that severe and extensive forest modification has occurred across all biogeographic regions of the world. Consequently, indices only using forest extent may inadequately capture the true impact of human activities on forests, and are insensitive to many drivers of forest modification and the resulting losses of forest benefits.

A plan is clearly needed to put in place retention strategies for the remaining forests with high integrity, tailored towards the context in each country or jurisdiction and its different forest types ^32,33^, because such areas are known to hold exceptional value. Avoiding the loss of integrity is a better strategy than aiming to restore forest condition after it is lost, because restoration is more costly, has a risk of failure, and is unlikely to lead to full recovery of benefits ^5^. For the forests with highest integrity to be retained they should ideally be mapped using nationally appropriate criteria by the countries that hold them, formally recognized, prioritized in spatial plans, and placed under effective management (e.g. protected areas and other effective conservation areas, lands under Indigenous control etc.). These forests must be protected from industrial development impacts that degrade them through sensible public and private sector policy that is effective at relevant scales ^12,34^. Our global assessment reveals where these places are found, and can be refined at more local scales where better data are available.

Around a third of global forests had already been cleared by 2000 ^35^, and we show that at least 59% of what remains has low to medium integrity, with > 50% falling in these two broad categories in every biogeographical realm. These levels of human modification result partly from the large areas affected by relatively diffuse anthropogenic pressures whose presence is inferred near forest edges, and by lost connectivity. We also map a surprising level of more localized, observed pressures, such as infrastructure and recent forest loss, which are seen in nearly a third of forested pixels worldwide.

Conservation strategies in these more heavily human-modified forests should focus on securing any remaining fragments of forests in good condition, proactively protecting those forests most vulnerable to further modification ^7^ and planning where restoration efforts might be most effective ^36-38^. In addition, effective management of production forests is needed to sustain yields without further worsening their ecological integrity ^39^. More research is required on how to prioritize, manage, and restore forests with low to medium integrity ^38,40^, and the FLII presented here might prove useful for this, for example, by helping prioritize where the best returns on investment are, in combination with other sources of data ^41^.

Loss of forest integrity severely compromises many benefits of forests that are central to achieving multiple Sustainable Development Goals and other societal targets ^42,43^. Therefore, governments must adopt policies and strategies to retain and restore the ecological integrity of their forests, whilst ensuring that the solutions are also economically viable, socially equitable, and politically acceptable within complex and highly diverse local contexts. This is an enormous challenge and our efforts to map the degree of forest modification are designed both to raise awareness of the importance of the issue, and to support implementation through target setting, evidence-based planning, and enhanced monitoring efforts.

Whilst policy targets for halting deforestation are generally precise and ambitious, only vague targets are typically stipulated around reducing levels of forest modification ^9,44^. We urgently need SMART (specific, measurable, achievable, realistic, and time-bound) goals and targets for maintaining and restoring forest integrity that directly feed into higher-level biodiversity, climate, land degradation, and sustainable development goals ^45^. These types of targets should be included within an over-arching target on ecosystems within the post-2020 Global Biodiversity Framework, which is currently being negotiated among Parties to the CBD ^46^. This target should be outcome-focused and address both the extent and the integrity of ecosystems (e.g. using FLII for forests), in a way that enables quantitative, measurable goals to be set but allows flexibility for implementation between Parties.

In addition to broader goals in global frameworks, the retention and restoration of forest integrity should also be addressed in nationally-defined goals embodied in, and aligned between, Nationally Determined Contributions under the UNFCCC, efforts to stop land degradation and achieve land degradation neutrality under the UNCCD, and National Biodiversity Strategy and Action Plans under the CBD. Since no single metric can capture all aspects of a country’s environmental values, efforts to conserve high levels of forest integrity should be complemented by consideration of areas support important values according to other measures (e.g. Key Biodiversity Areas ^47^ and notable socio-cultural landscapes).

The overall level and pervasiveness of impacts on Earth’s remaining forests is likely even more severe than our findings suggest, because some input data layers, despite being the most comprehensive available, are still incomplete as there are lags between increases in human pressures and our ability to capture them in spatial datasets e.g., infrastructure, ^48,49, see also Fig. S1 and text S5^. For example, roads and seismic lines used for natural resource exploration and extraction in British Columbia, Canada, are not yet fully reflected in global geospatial datasets Fig. S1; see also ^50^. Furthermore, because natural fires are such an important part of the ecology of many forest systems (e.g. boreal forests) and because we cannot consistently identify anthropogenic fires from natural fires at a global scales ^51^ we have taken a strongly conservative approach to fire in our calculations, treating all tree cover loss in 10 km pixels where fire was the dominant driver *(23)* as temporary, and not treating such canopy loss as evidence of observed human pressure. Varying these assumptions where human activity is shown to be causing permanent tree cover losses, increasing fire return frequencies, or causing fire in previously fire-free systems would result in lower forest extent and/or lower forest integrity scores in some regions than we report.

We map forest integrity based on quantifiable processes over the recent past (since 2000). In some areas modification that occurred prior to this (e.g. historical logging) is not detectable by our methods but may have influenced the present-day integrity of the forest so, in such cases, we may overestimate forest integrity. This is another reason why our index should be considered as conservative, and we therefore recommend that the index be used alongside other lines of evidence to determine the absolute level of ecological integrity of a given area. Moreover, the definition of forest in this study is all woody vegetation taller than 5 m, following ^22^ and hence includes not only naturally regenerated forests but also tree crops, planted forests, wooded agroforests and urban tree cover in some cases. Users should be mindful of this when interpreting the results, especially when observing areas with low forest integrity scores. Inspection of the results for selected countries with reliable plantation maps ^52^ shows that the great majority of planted forests have low forest integrity scores, because they are invariably associated with dense infrastructure, frequent canopy replacement and patches of farmland.

We note our measure of forest integrity does not address past, current and future climate change. As climate change affects forest condition both directly and indirectly, this is a clear shortfall and needs research attention. The same is true for invasive species, as there is no globally coherent data on the range of those invasive species that degrade forest ecosystems, although this issue is indirectly addressed since the presence of many invasive species are likely spatially correlated with the human pressures that we use as drivers in our model ^53^. If global data became available it would also be valuable to incorporate governance effectiveness into our model, because there are potentially contexts (e.g. well-managed protected areas and community lands, production forests under ‘sustainable forest management’) where the impacts associated with the human pressures we base our map on are at least partially ameliorated ^39^, and enhanced governance is also likely to be a significant component of some future strategies to maintain and enhance forest integrity.

The framework we present has great potential to be tailored for use at smaller scales, ranging from regional to national and sub-national scales, and even to individual management units. Forest definitions and the relative weights of the global parameters we use can be adjusted to fit local contexts and, in many cases, better local data could be substituted, or additional variables incorporated. This would increase the precision of the index in representing local realities, and increase the degree of ownership amongst national and local stakeholders whose decisions are so important in determining forest management trajectories.

## Methods

To produce our global Forest Landscape Integrity Index (FLII), we combined four sets of spatially explicit datasets representing: (i) forest extent ^22^; (ii) ‘observed’ pressure from high impact, localized human activities for which spatial datasets exist, specifically: infrastructure, agriculture, and recent deforestation ^53^; (iii) ‘inferred’ pressure associated with edge effects ^54^, and other diffuse processes, (e.g. activities such as hunting and selective logging) ^55^ modelled using proximity to observed pressures; and iv) anthropogenic changes in forest connectivity due to forest loss ^56 see Table S1 for data sources^. These datasets were combined to produce an index score for each forest pixel (300m), with the highest scores reflecting the highest forest integrity (Fig 1), and applied to forest extent for the start of 2019. We use globally consistent parameters for all elements (i.e. parameters do not vary geographically). All calculations were conducted in Google Earth Engine (GEE) ^57^.

### Forest extent

We derived a global forest extent map for 2019 by subtracting from the Global Tree Cover product for 2000 ^22^ annual Tree Cover Loss 2001-2018, except for losses categorized by Curtis and colleagues ^23^ as those likely to be temporary in nature (i.e. those due to fire, shifting cultivation and rotational forestry). We applied a canopy threshold of 20% based on related studies e.g. ^29,58^ and resampled to 300m resolution and used this resolution as the basis for the rest of the analysis (see text S1 for further mapping methods).

### Observed human pressures

We quantify observed human pressures (P) within a pixel as the weighted sum of impact of infrastructure (I; representing the combined effect of 41 types of infrastructure weighted by their estimated general relative impact on forests (Table S3), agriculture (A) weighted by crop intensity (indicated by irrigation levels), and recent deforestation over the past 18 years (H; excluding deforestation from fire, see Discussion). Specifically, for pixel i:

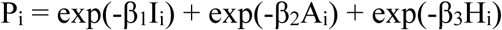

whereby the values of β were selected so that the median of the non-zero values for each component was 0.75. This use of exponents is a way of scaling variables with non-commensurate units so that they can be combined numerically, while also ensuring that the measure of observed pressure is sensitive to change (increase or decrease) in the magnitude of any of the three components, even at large values of I, A or H. This is an adaptation of the ‘Human Footprint’ methodology ^53^. See text S3 for further details.

### Inferred human pressures

Inferred pressures are the diffuse effects of a set of processes for which directly observed datasets do not exist, that include microclimate and species interactions relating to the creation of forest edges ^59^ and a variety of intermittent or transient anthropogenic pressures such as: selective logging, fuelwood collection, hunting; spread of fires and invasive species, pollution, and livestock grazing ^55,60,61^. We modelled the collective, cumulative impacts of these inferred effects through their spatial association with observed human pressure in nearby pixels, including a decline in effect intensity according to distance, and a partitioning into stronger short-range and weaker long-range effects. The inferred pressure (P’) on pixel *i* from source pixel *j* is:

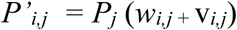

where w_*i,j*_ is the weighting given to the modification arising from short-range pressure, as a function of distance from the source pixel, and v_*i,j*_ is the weighting given to the modification arising from long-range pressures.

Short-range effects include most of the processes listed above, which together potentially affect most biophysical features of a forest, and predominate over shorter distances. In our model they decline exponentially, approach zero at 3 km, and are truncated to zero at 5 km (see text S4).

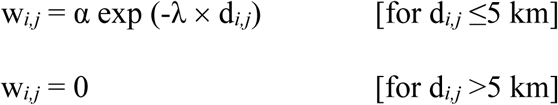

where α is a constant set to ensure that the sum of the weights across all pixels in range is 1.85 (see below), λ is a decay constant set to a value of 1 (see ^62^ and other references in text S4) and d_*i,j*_ is the Euclidean distance between the centres of pixels *i* and *j* expressed in units of km.

Long-range effects include over-exploitation of high socio-economic value animals and plants, changes to migration and ranging patterns, and scattered fire and pollution events. We modelled long-range effects at a uniform level at all distances below 6 km and they then decline linearly with distance, conservatively reaching zero at a radius of 12 km ^55,63 and other references in text S4^:

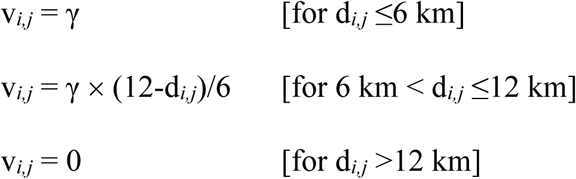

Where γ is a constant set to ensure that the sum of the weights across all pixels in range is 0.15 and d_*i,j*_ is the Euclidean distance between the centres of pixels *i* and *j*, expressed in kilometres.

The form of the weighting functions for short- and long-range effects and the sum of the weights (α+γ) were specified based on a hypothetical reference scenario where a straight forest edge is adjacent to a large area with uniform human pressure, and ensuring that in this case total inferred pressure immediately inside the forest edge is equal to the pressure immediately outside, before declining with distance. γ is set to 0.15 to ensure that the long-range effects conservatively contribute no more than 5% to the final index in the same scenario, based on expert opinion and supported e.g. Berzaghi *et al.* ^64^ regarding the approximate level of impact on values that would be affected by severe defaunation and other long-range effects.

The aggregate effect from inferred pressures (P’) on pixel *i* from all *n* pixels within range (*j=1* to *j=n*) is then the sum of these individual, normalized, distance-weighted pressures, i.e.

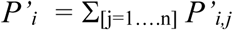

### Loss of forest connectivity

Average connectivity of forest around a pixel was quantified using a method adapted from Beyer *et al.* ^56^. The connectivity C*i* around pixel *i* surrounded by n other pixels within the maximum radius (numbered *j*=1, 2…n) is given by:

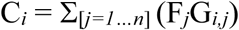

where F*j* is the forest extent is a binary variable indicating if forested (1) or not (0) and G*i,j* is the weight assigned to the distance between pixels *i* and *j*. G*i,j* uses a normalized Gaussian curve, with σ = 20km and distribution truncated to zero at 4σ for computational convenience (see text S3). The large value of σ captures landscape connectivity patterns operating at a broader scale than processes captured by other data layers. C_*i*_ ranges from 0 to 1 (C_*i*_ ∈ [0,1]).

Current Configuration (CC_*i*_) of forest extent in pixel i was calculated using the final forest extent map and compared to the Potential Configuration (PC) of forest extent without extensive human modification, so that areas with naturally low connectivity, e.g. coasts and natural vegetation mosaics, are not penalized. PC was calculated from a modified version of the map of Laestadius *et al.* ^35^ and resampled to 300 m resolution (see text S2 for details). Using these two measures, we calculated Lost Forest Configuration (LFC) for every pixel as:

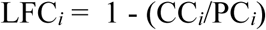

Values of CC_*i*_/PC_*i*_ >1 are assigned a value of 1 to ensure that LFC is not sensitive to apparent increases in forest connectivity due to inaccuracy in estimated potential forest extent – low values represent least loss, high values greatest loss (LFC_*i*_ ∈ [0,1]).

### Calculating the Forest Landscape Integrity Index

The three constituent metrics, LFC, P and P’, all represent increasingly modified conditions the larger their values become. To calculate a forest integrity index in which larger values represent less degraded conditions we therefore subtract the sum of those components from a fixed large value (here, 3). Three was selected as our assessment indicates that values of LFC + P + P’ of 3 or more correspond to the most severely degraded areas. The metric is also rescaled to a convenient scale (0-10) by multiplying by an arbitrary constant (10/3). The FLII for forest pixel *i* is thus calculated as:

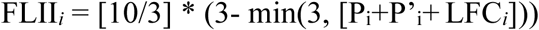

where FLII_*i*_ ranges from 0 - 10, forest areas with no modification detectable using our methods scoring 10 and those with the most scoring 0.

### Illustrative forest integrity classes

Whilst a key strength of the index is its continuous nature, the results can also be categorized for a range of purposes. In this paper three illustrative classes were defined, mapped and summarized to give an overview of broad patterns of integrity in the world’s forests. The three categories were defined as follows.

#### *High Forest Integrity* (scores ≥9.6)

Interiors and natural edges of more or less unmodified naturally-regenerated (i.e. non-planted) forest ecosystems, comprised entirely or almost entirely of native species, occurring over large areas either as continuous blocks or natural mosaics with non-forest vegetation; typically little human use other than low intensity recreation or spiritual uses and/or low intensity extraction of plant and animal products and/or very sparse presence of infrastructure; key ecosystem functions such as carbon storage, biodiversity and watershed protection and resilience expected to be very close to natural levels (excluding any effects from climate change) although some declines possible in the most sensitive elements (e.g. some high value hunted species).

#### *Medium Forest Integrity* (scores >6.0 but <9.6)

Interiors and natural edges of naturally-regenerated forest ecosystems in blocks smaller than their natural extent but large enough to have some core areas free from strong anthropogenic edge effects (e.g. set asides within forestry areas, fragmented protected areas), dominated by native species but substantially modified by humans through a diversity of processes that could include fragmentation, creation of edges and proximity to infrastructure, moderate or high levels of extraction of plant and animal products, significant timber removals, scattered stand-replacement events such as swidden and/or moderate changes to fire and hydrological regimes; key ecosystem functions such as carbon storage, biodiversity, watershed protection and resilience expected to be somewhat below natural levels (excluding any effects from climate change).

#### *Low Forest Integrity* (score ≤6.0)

Diverse range of heavily modified and often internally fragmented ecosystems dominated by trees, including (i) naturally regenerated forests, either in the interior of blocks or at edges, that have experienced multiple strong human pressures, which may include frequent stand-replacing events, sufficient to greatly simplify the structure and species composition and possibly result in significant presence of non-native species, (ii) tree plantations and, (iii) agroforests; in all cases key ecosystem functions such as carbon storage, biodiversity, watershed protection and resilience expected to be well below natural levels (excluding any effects from climate change).

The numerical category boundaries were derived by inspecting FLII scores for a wide selection of benchmark locations whose forest integrity according to the category definitions was known to the authors, see text S6 and Table S4.

## Protected areas analysis

Data on protected area location, boundary, and year of inscription were obtained from the February 2018 World Database on Protected Areas ^65^. Following similar global studies e.g. ^66^, we extracted protected areas from the WDPA database by selecting those areas that have a status of “designated”, “inscribed”, or “established”, and were not designated as UNESCO Man and Biosphere Reserves. We included only protected areas with detailed geographic information in the database, excluding those represented as a point only. To assess integrity of protected forest, we extracted all 300m forest pixels that were at least 50% covered by a formal protected area and measured the average FLII score.

## Acknowledgments

We thank Peter Potapov, Dmitry Aksenov, and Matthew Hansen for comments and advice.

## Author contributions

Conceived and design the study: HG, TE and JEW, collected data and developed the model: AD, HG, TE, HB, RS, analyzed and interpreted the results: AD, HG, TE, HB, RS, KJ, JEW, wrote draft manuscript: HG, TE and JEW, contributed to the writing of the manuscript: all co-authors.

## Data and materials availability

Data will be available for download when published.

## Supplementary Materials

### Text S1. Mapping forest extent

We generated a preliminary base map of global forest extent for the start of 2019 at 30 m resolution by subtracting annual Tree Cover Loss 2001-2018 (with exceptions noted in the next paragraph) from the Global Tree Cover 2000 product ^22^ using a canopy cover threshold of 20%. This is one of the most widely used tree cover datasets globally, so it has been widely tested in many settings and its strengths and constraints are well understood. It has many advantages, including its high resolution, high accuracy, global coverage, annual time series and good prospects of sustainability in the coming years. The definition of forest in the source dataset is all woody vegetation taller than 5 m and hence includes naturally regenerated forests as well as tree crops, planted forests, wooded agroforests and urban tree cover. No globally consistent dataset was available that allowed natural and planted tree cover to be consistently distinguished in this study. Therefore, we should be mindful of the many differences between planted and natural tree cover (e.g.^67^).

More than 70% of the tree cover loss shown by the Hansen *et al.* ^22^ products has been found to be in 10 km pixels where the dominant loss driver is temporary and so tree cover is expected to return above the forest definition threshold within a short period ^23^. It is important to take account of this issue as treating all such areas as permanent loss would severely under-estimate current forest cover in many regions. However, no global map of forest cover gain exists for the study period other than the 2000-2012 gain product from Hansen *et al.* ^22^, so we developed an alternative approach. When removing annual loss shown by the Global Tree Cover Loss product cited above we elected not to remove any loss that was in a 10 km pixel categorized by Curtis *et al.* ^23^ as dominated by temporary loss under the categories of fire, shifting cultivation or rotational forestry. This resulted in the adjusted preliminary forest base map. The balance of evidence is that the great majority of such areas would have begun to regenerate and hence qualify as forest by our definition again by 2019 or soon after ^23^. The anthropogenically disturbed nature of many of these areas of temporary tree cover loss and recovery is reflected in scoring within the index, because temporary tree cover loss in the categories of shifting cultivation or rotational forestry is treated as an observed pressure. We do not treat tree cover loss through fire as an observed pressure, because fires are often part of natural processes, especially in the boreal zone. This makes our global index conservative as a measure of degradation in these zones, because in some locations fires are anthropogenic in nature.

The adjusted preliminary base map was then resampled to a final base map for 2019 at 300m resolution using a pyramid-by-mode decision rule, with the resulting pixels simply classified as forest or non-forest based on a majority rule. The FLII was calculated for every forest pixel but not for non-forest pixels. GEE performs calculations in WGS84. Supplementary analyses outside GEE were applied using a Mollweide equal-area projection.

### Text S2. Mapping potential forest configuration

Potential connectivity (PC) is calculated from an estimate of the potential extent of the forest zone taken from Laestadius *et al.* ^35^, treating areas below 25% crown cover (this was the nearest class to the threshold used in our tree cover dataset of 20%) as non-forest and resampling to 300 m resolution. To minimize false instances of lost connectivity and ensure measures of forest modification are conservative we masked from this data layer areas which we believe to include a significant proportion of naturally unforested land using selected land-cover categories in ESA (^68^; see Table S1). Because these natural non-forest patches are shown in the Hansen *et al.* ^22^ dataset but not Laestadius *et al.* ^35^, not excluding such classes would result in an inflated estimate of the loss of connectivity and hence the level of degradation. We have elected to remain conservative in our estimate of modification.

### Text S3. Mapping observed human pressure

Several recent analyses have developed composite, multi-criteria indices of human pressure to provide assessments of ecosystem condition for the USA ^69^ or globally ^26,70,71^. Thompson *et al.* ^72^ set out a framework specific to forest ecosystems that could indicate modification through a balanced mix of available pressure and state variables. We adapted the methodology of Venter et al. ^26^, informed by the other studies cited, to generate measures of (i) the modification of forest associated with observed human pressure from infrastructure, agriculture and deforestation and (ii) the more diffuse inferred modification effects (e.g. edge effects) whose presence is inferred from proximity to these focal areas of human activity. Edge effects resulting entirely from natural processes are excluded, because they do not represent modification by our definition, although, like many other natural factors, they do also have a role in determining ecosystem benefits.

#### Infrastructure

We generated the infrastructure (I’) data layer by rasterizing the OpenStreetMap data ^73^ from Feb 2018, using weights for each type of infrastructure as noted in Table S3. The weights were derived from authors’ expert opinion and experimentation with weights according to their relative impact on forest condition.

#### Agriculture

For agriculture (A’) we made a global binary composite of the croplands datasets produced by the USGS (Table 1) at 30 m resolution, and weighted each cropped pixel at this resolution by the likely intensity of cropping using the global irrigation dataset at 1km resolution (Teluguntla et al, ^74^), with values of Irrigation Major = 2, Irrigation Minor = 1.5, Rainfed = 1. The average cropping intensity (including uncropped areas, which score zero) was then calculated across the whole of each 300 m pixel of our final basemap.

#### Deforestation

For deforestation (H’) we made a binary composite of tree cover loss 2001-2018 at 30 m resolution ^22^, masked out 30 m pixels already classified as agriculture in the preceding step to avoid double-counting, and excluded loss predicted by Curtis *et al*. ^23^ to be most likely caused by fires, to give a conservative data layer of recent permanent and temporary tree cover loss indicative of human activity in the immediate vicinity. We excluded small clusters of 6 or fewer pixels (0.54 ha) because they may have been natural tree cover loss (e.g. small windthrows) or classification errors. Each 30 m pixel was then weighted by its year of loss, giving higher weight to the most recent loss (2001 = 1, 2002 = 2, etc.). The average ‘recentness’ of deforestation (including areas not deforested, which score zero) was then calculated across the whole of each 300 m pixel of our base map.

#### Transformations

The exponential transformations described in the main text were used to convert I’, A’ and H’ to the variables I, A and H respectively.

### Text S4. Modelling inferred pressures using proximity to observed pressures

Each cell also experiences modification as a result of pressures originating from nearby cells that have observed human pressures, largely through the family of processes known as ‘edge effects’^54^. Edge effects are partly a result of the changes relating to biophysical factors (such as humidity, wind, temperature and the increased presence of non-forest species) that accompany the creation of new edges in formerly continuous forest (as exemplified by the carefully controlled studies in tropical forests summarized by Laurance *et al.* ^59^). They also result in part from the increased pressure associated with human activities within tropical forest near to edges such as logging ^61^, anthropogenic fire ^60^, hunting ^55^, livestock grazing, pollution, visual and auditory disturbances, etc. These multiple factors are synergistic and so we model them together, notwithstanding regional and local variations in the relative intensity of each one.

We model the inferred effect caused by each nearby source cell as a function of (a) the observed human pressure observed in that source cell and (b) a decline in the intensity of edge effects with distance from the source cell, based on a review of the literature. We then determine the total inferred effect on a given cell by summing the individual effects from all source cells within a certain range.

Two complementary types of inferred effect are modelled and added together. One relates to the diverse, strong, relatively short-range edge effects which decay to near zero over a few kilometers and have the potential to affect most biophysical features of a forest to a greater or lesser extent.

The other relates to weaker, longer-range effects such as over-hunting of high-value animals that affect fewer biophysical features of a forest (and so have a much smaller maximum effect on overall integrity) but can nonetheless have detectable effects in locations more than 10 km from the nearest permanent human presence.

The literature on the spatial influence of short-term effects uses a variety of mathematical descriptors, in two broad categories – continuous variables and distance belts. As we wish to model edge effects as a continuous variable we concentrated on studies that have taken a similar approach, and used distance-belt studies as ancillary data.

Chaplin-Kramer *et al.* ^62^ is a good example of a continuous variable approach, estimating detailed biomass loss curves near tropical forest edges. Because they analyze a key forest condition variable with a very large pantropical dataset we hypothesize that the exponential declines in degradation with distance that they find are likely to be a common pattern and so we use a similar framework for our more general model of degradation. We consider that a model of exponential decay is also a sufficient approximation to the evidence presented by some authors as graphs without an associated mathematical model (e.g., ^60,75^) or analyzed using logistic regression (e.g., ^76^). In our model we set the exponential decay constant to be broadly consistent with these four studies, resulting in degradation at 1 km inside a forest that is approximately 37% of that at the forest edge, declining to 14% at 2 km and near zero at 3 km. We truncate the distribution at 5 km to minimize computational demands.

Distance-belt studies define the width of a belt within which edge effects are considered to occur, and beyond which forests are considered to be free of edge effect. Belts of 1 km are commonly used (e.g., ^54^) but smaller distances may be used for specific parameters (e.g. 300 m for biomass reduction near edges in DRC’s primary forests; ^27^). Our continuous variable approach is broadly consistent with these studies, with the majority of our modelled degradation within a 1 km belt and little extending beyond 2 km. While most individual edge effects reported in the literature penetrate less than 100-300 m (e.g., ^59,77^) most of the effects reported on in these studies relate to the changed natural factors mentioned in an earlier paragraph, and are likely to be dwarfed in both intensity and extent by edge effects relating to spillovers of human activity, so our model emphasizes the spatial distribution of the latter (e.g., ^60^). We consider our model of the levels of modification to be conservative.

For the weaker, more widespread long-range effects we use recent large-scale studies of defaunation, which is one of the key long-range pressures and also acts as a proxy for other threats including harvest of high value plants (such as eaglewood *Aquilaria* spp. in tropical Asia), occasional remote fires, pollution associated with artisanal mining, etc. We adopt a simplified version of the distribution used by Peres *et al.* ^55^ to model hunting around settlements in the Amazon, which sets 2σ=12 km; this is likely conservative compared to evidence for hunting-related declines in forest elephants in central Africa up to 60 km from roads ^63^ and the extensive declines in large-bodied quarry species in remote areas in many regions modelled by Benitez-Lopez *et al.* ^78^.

### Text S5. Limitations in data: example with infrastructure data in British Columbia, Canada

OpenStreetMap (OSM) represents the most detailed publicly available relevant global dataset but is nonetheless noted to be incomplete, even for one of the most heavily used categories of infrastructure, paved roads ^48^. No global assessment is available for the completeness of other categories in the dataset. One of the key categories for forest integrity, unpaved roads used for resource extraction, has been shown to be incomplete over much of insular South-east Asia ^49^. In Canada, for example, roads and other linear corridors used to explore, access and extract natural resources (e.g., logging, oil and gas, and minerals) are sometimes missing. Government data for the province of British Columbia (available at https://catalogue.data.gov.bc.ca/dataset/digital-road-atlas-dra-master-partially-attributed-roads) demonstrates, for example, the larger extent and density of regional roads as compared to OSM (Fig S1).

### Text S6. Classification of Forest Landscape Integrity Index scores

In this paper, three illustrative classes were defined, mapped and summarized to give an overview of broad patterns of degradation in the world’s forests. Three categories were defined as set out in the Materials and Methods. To determine the approximate levels of the FLII associated with these three categories, benchmark locations were selected in sites that could unambiguously be assigned to one of the categories using the authors’ personal knowledge. At each site a single example pixel was selected within a part of the area with relatively uniform scores. The sample points are summarized in Table S4; they are widely spread across the world to ensure that the results are not only applicable to a limited region. The scores at these points suggest the following category boundaries:

- High FLII – 9.6-10
- Medium FLII – 6-9.6
- Low FLII – 0-6

**Table S1.**
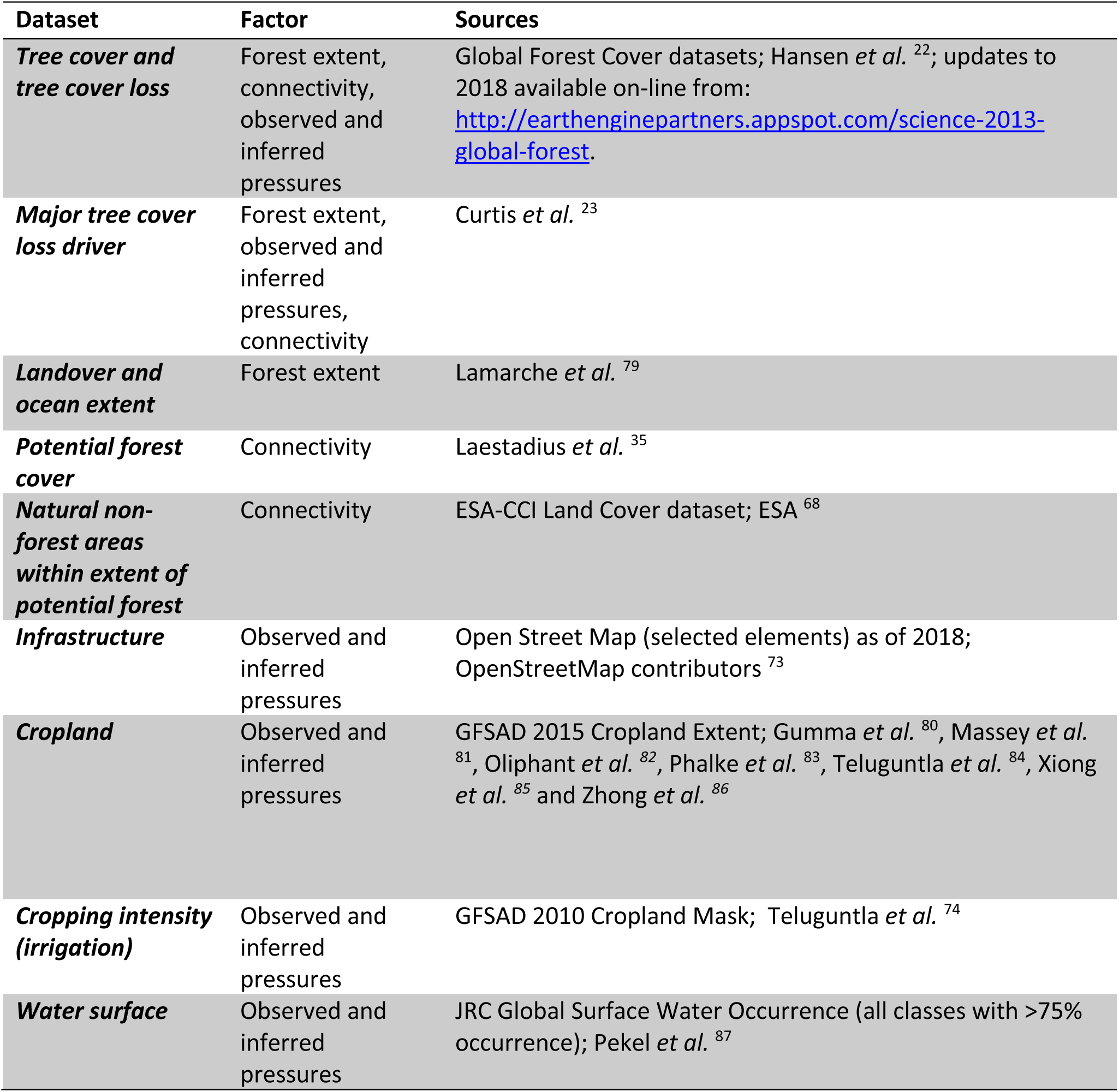
The datasets used to develop the Forest Ecosystem Integrity Index. The factor column indicates the component of the index the dataset was used in.

**Table S2.**
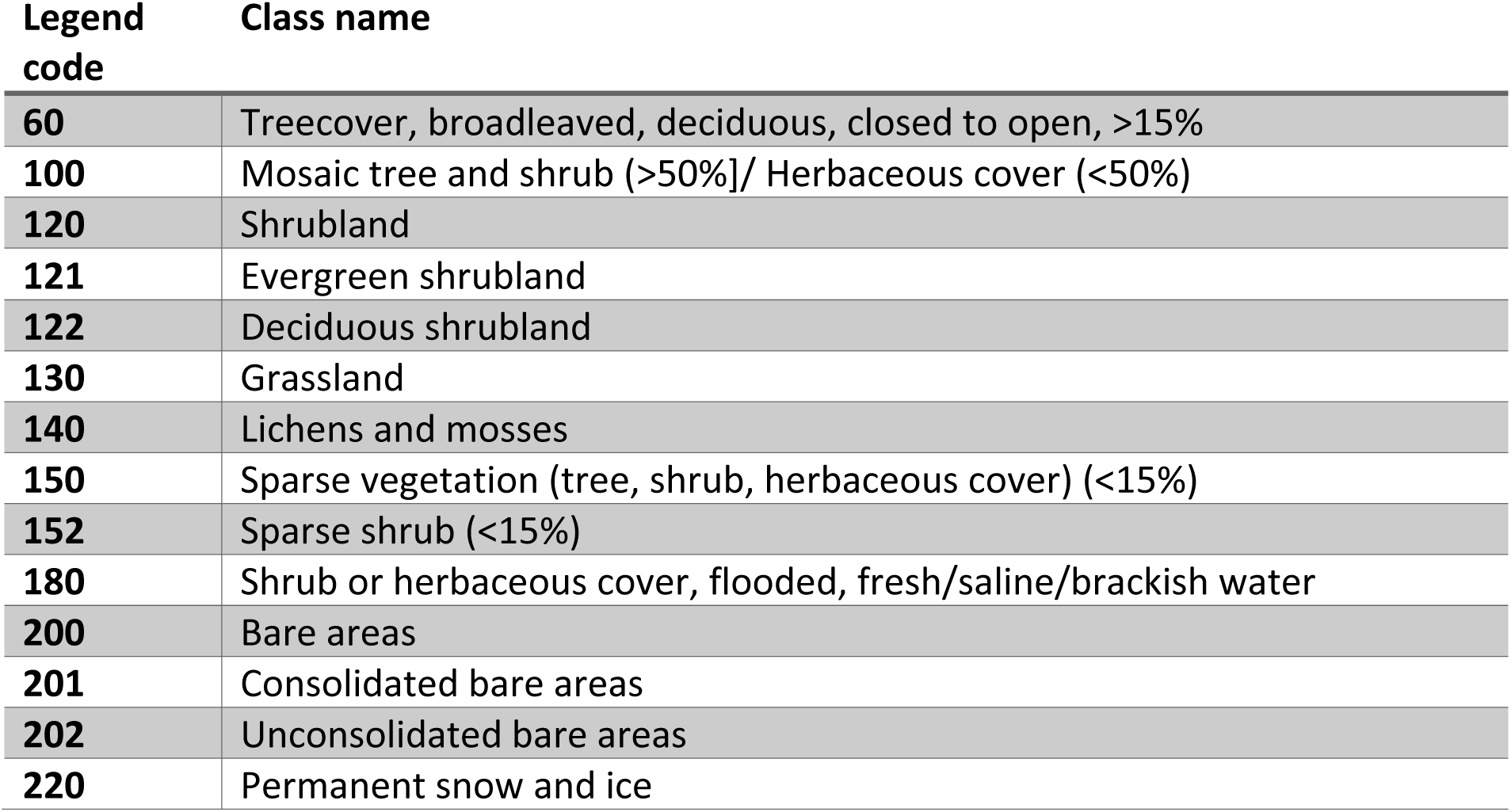
Classes in ESA-CCI dataset excluded from our potential forest cover layer because they overlap extensively with potential forest cover mapped by Laestadius *et al.* ^35^ but contain significant areas of natural non forest

**Table S3.**
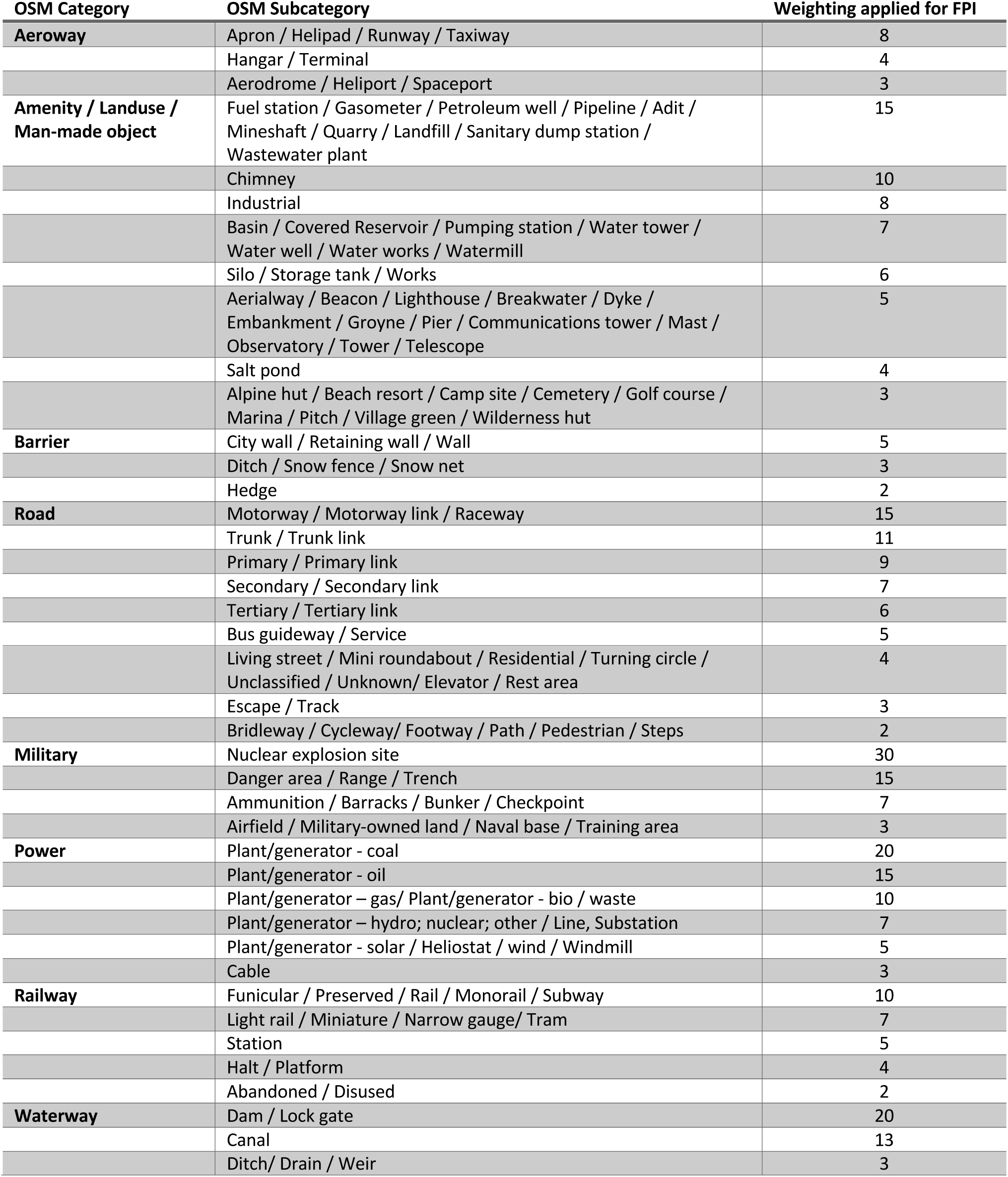
Weightings used for Open Street Map (OSM) to combine into the Infrastructure data layer.

**Table S4.** Points assessed to determine category boundaries for classifying the FHI into high, medium and low classes.

| Category | Code | Point description | Country | Point Score |
| --- | --- | --- | --- | --- |
| High | 103 | Interior of Lopé National Park | Gabon | 10.000 |
| High | 106 | Interior of Tai National Park | Cote d'Ivoire | 10.000 |
| High | 108 | Interior of Pacaya-Samiria National Reserve | Peru | 10.000 |
| High | 109 | Interior of Central Suriname Nature Reserve | Suriname | 10.000 |
| High | 116 | Interior of Liard River area | Canada | 10.000 |
| High | 101 | Interior of Okapi Faunal Reserve | DRC | 9.997 |
| High | 104 | Interior of Nyungwe National Park | Rwanda | 9.992 |
| High | 111 | Interior of Rio Platano Biosphere Reserve | Honduras | 9.990 |
| High | 102 | Interior of Odzala National Park | RoC | 9.974 |
| High | 117 | Interior of Wells Gray Provincial Park | Canada | 9.972 |
| High | 119 | Interior of Øvre Pasvik National Park | Norway | 9.944 |
| High | 115 | Interior of Tasmania Wilderness World Heritage Area | Australia | 9.918 |
| High | 107 | Interior of Marojejy National Park | Madagascar | 9.910 |
| High | 112 | Interior of Khao Yai National Park | Thailand | 9.908 |
| High | 105 | Interior of Niassa Special Reserve | Mozambique | 9.819 |
| High | 110 | Interior of Maya Biosphere Reserve | Guatemala | 9.798 |
| High | 114 | Interior of Batang Ai National Park | Malaysia | 9.756 |
| High | 118 | Interior of Quetico Provincial Park | Canada | 9.750 |
| High | 113 | Interior of Sundarbans National Park | Bangladesh | 9.606 |
| Medium | 215 | Interior of Bialowieza National Park | Poland | 9.086 |
| Medium | 208 | Interior of Mabira Central Forest Reserve | Uganda | 9.067 |
| Medium | 211 | Area of selective logging | Gabon | 8.840 |
| Medium | 219 | Near main tourism corridor, Mt Myohyang National Park | DPR Korea | 8.762 |
| Medium | 203 | Interior of Phnom Kulen Wildlife Sanctuary | Cambodia | 8.710 |
| Medium | 210 | Area of selective logging | Guyana | 8.364 |
| Medium | 202 | Interior of Dong Hua Sao National Protected Area | Lao PDR | 8.078 |
| Medium | 212 | Area of selective logging | DRC | 7.981 |
| Medium | 206 | Interior of Manga Forest Reserve | Tanzania | 7.960 |
| Medium | 207 | Near margin of Nyungwe National Park | Rwanda | 7.938 |
| Medium | 204 | South part of Nagarhole National Park | India | 7.759 |
| Medium | 213 | Area of selective logging | Cameroon | 7.379 |
| Medium | 201 | Tat Leuk, Phou Khaokhoay National Protected Area | Lao PDR | 7.251 |
| Medium | 216 | Interior of Loch Garten Nature Reserve | UK | 7.146 |
| Medium | 209 | Area of selective logging | Congo | 6.734 |
| Medium | 217 | Tourism area, Lamington National Park | Australia | 6.729 |
| Medium | 214 | Lowlands of Guanacaste National Park | Costa Rica | 6.719 |
| Medium | 218 | Near margin of Sepilok Forest Reserve | Malaysia | 6.353 |
| Medium | 205 | Interior of Similajau National Park | Malaysia | 6.130 |
| Low | 305 | Dong Nathat | Lao PDR | 5.638 |
| Low | 317 | Foothills of Mt Makiling | Philippines | 5.395 |
| Low | 310 | Suburban woodlot, Dobbs Ferry | USA | 4.710 |
| Low | 309 | Jozani Forest Reserve | Tanzania | 4.680 |
| Low | 316 | Foothills of Mt Canlaon | Philippines | 4.597 |
| **Low** | 320 | Forest fragment near Paramaribo | Suriname | 4.566 |
| **Low** | 302 | Central Park, New York | USA | 3.575 |
| **Low** | 301 | Bagley Wood, Oxford | UK | 3.525 |
| **Low** | 307 | Boeng Yeak Lom Protected Area | Cambodia | 3.323 |
| **Low** | 304 | Angkor Thom | Cambodia | 3.122 |
| **Low** | 315 | Forest in rural complex, Mambasa area | DRC | 2.689 |
| **Low** | 312 | Woodland in Beaumont area | USA | 2.581 |
| **Low** | 318 | Swidden near Andoung Kraloeng village | Cambodia | 2.304 |
| **Low** | 319 | Forest mosaic near Kaev Seima village | Cambodia | 2.187 |
| **Low** | 303 | Thetford Forest | UK | 2.082 |
| **Low** | 313 | Woodland in Augusta area | USA | 0.686 |
| **Low** | 314 | Woodland in Emporia area | USA | 0.589 |
| **Low** | 311 | River Park, Chicago | USA | 0.566 |
| **Low** | 306 | Houei Nhang Forest Reserve | Lao PDR | 0.000 |
| **Low** | 308 | Pugu Forest Reserve | Tanzania | 0.000 |

**Table S5.** Mean Forest Landscape Integrity Index scores and areas for forest integrity categories by country.

| Country | Mean<br>FHI | Low health<br>(km<sup>2</sup>) | Medium health<br>(km<sup>2</sup>) | High health<br>(km<sup>2</sup>) | Total forest<br>area (km<sup>2</sup>) |
| --- | --- | --- | --- | --- | --- |
| **Afghanistan** | 8.85 | 90 | 1,475 | 977 | 2,542 |
| **Albania** | 6.77 | 2,426 | 5,256 | 122 | 7,805 |
| **Algeria** | 5.22 | 7,418 | 6,044 | 81 | 13,543 |
| **Andorra** | 4.45 | 170 | 49 | 0 | 219 |
| **Angola** | 8.35 | 105,487 | 284,054 | 315,895 | 705,436 |
| **Antigua and Barbuda** | 4.72 | 114 | 92 | 0 | 206 |
| **Argentina** | 7.21 | 98,249 | 189,966 | 72,557 | 360,772 |
| **Armenia** | 5.46 | 1,894 | 1,681 | 3 | 3,577 |
| **Australia** | 7.22 | 117,672 | 239,624 | 103,852 | 461,148 |
| **Austria** | 3.55 | 36,666 | 12,422 | 21 | 49,109 |
| **Azerbaijan** | 6.55 | 4,820 | 7,189 | 1,534 | 13,543 |
| **Bahamas** | 7.35 | 741 | 1,935 | 399 | 3,075 |
| **Bangladesh** | 5.45 | 10,013 | 7,251 | 1,947 | 19,211 |
| **Belarus** | 3.63 | 77,870 | 20,847 | 91 | 98,808 |
| **Belgium** | 1.36 | 8,803 | 297 | 0 | 9,099 |
| **Belize** | 6.15 | 7,004 | 7,957 | 2,744 | 17,705 |
| **Benin** | 5.86 | 4,724 | 3,698 | 1,769 | 10,191 |
| **Bhutan** | 8.85 | 1,620 | 16,769 | 10,140 | 28,529 |
| **Bolivia** | 8.47 | 78,745 | 280,532 | 272,007 | 631,284 |
| **Bosnia and**<br>**Herzegovina** | 5.99 | 13,387 | 17,031 | 574 | 30,993 |
| **Botswana** | 9.13 | 13 | 187 | 372 | 572 |
| **Brazil** | 7.52 | 1,374,902 | 1,354,961 | 2,338,101 | 5,067,963 |
| **Brunei Darussalam** | 7.71 | 1,102 | 2,842 | 1,498 | 5,442 |
| **Bulgaria** | 6.09 | 18,884 | 26,325 | 847 | 46,057 |
| **Burundi** | 4.5 | 6,882 | 3,841 | 46 | 10,769 |
| **Cabo Verde** | 6.37 | 27 | 38 | 0 | 65 |
| **Cambodia** | 6.31 | 30,143 | 31,939 | 16,349 | 78,431 |
| **Cameroon** | 8 | 66,191 | 181,336 | 119,263 | 366,789 |
| **Canada** | 8.99 | 480,206 | 1,027,386 | 2,968,268 | 4,475,860 |
| **Central African**<br>**Republic** | 9.28 | 30,161 | 139,350 | 379,097 | 548,608 |
| **Chad** | 6.18 | 5,261 | 6,016 | 1,910 | 13,187 |
| **Chile** | 7.37 | 56,849 | 41,971 | 93,537 | 192,357 |
| **China** | 7.14 | 533,800 | 974,431 | 301,051 | 1,809,282 |
| **Colombia** | 8.26 | 150,737 | 272,442 | 428,320 | 851,499 |
| **Comoros** | 7.69 | 284 | 1,149 | 82 | 1,515 |
| **Congo** | 8.89 | 24,512 | 124,215 | 158,184 | 306,911 |
| **Congo DRC** | 7.56 | 533,118 | 935,508 | 727,983 | 2,196,608 |
| **Costa Rica** | 4.65 | 27,164 | 12,838 | 4,164 | 44,167 |
| **Cote d'Ivoire** | 3.64 | 158,010 | 41,005 | 7,288 | 206,303 |
| **Croatia** | 4.92 | 15,732 | 10,522 | 379 | 26,633 |
| **Cuba** | 5.4 | 22,605 | 18,460 | 1,632 | 42,697 |
| **Cyprus** | 7.06 | 388 | 1,026 | 18 | 1,432 |
| **Czechia** | 1.71 | 32,161 | 1,611 | 0 | 33,772 |
| **Denmark** | 0.5 | 5,756 | 31 | 0 | 5,787 |
| **Dominica** | 1.06 | 531 | 2 | 0 | 533 |
| **Dominican Republic** | 4.19 | 19,890 | 9,364 | 518 | 29,772 |
| **Ecuador** | 7.66 | 48,822 | 77,585 | 73,492 | 199,900 |
| **Egypt** | 0.56 | 4,772 | 218 | 69 | 5,059 |
| **El Salvador** | 4.05 | 8,837 | 2,947 | 0 | 11,784 |
| **Equatorial Guinea** | 7.99 | 3,982 | 17,595 | 5,007 | 26,585 |
| **Estonia** | 3.05 | 24,473 | 4,832 | 52 | 29,358 |
| **Ethiopia** | 7.16 | 52,652 | 84,430 | 44,397 | 181,479 |
| **Fiji** | 8.35 | 1,753 | 10,802 | 3,594 | 16,148 |
| **Finland** | 5.08 | 144,310 | 83,572 | 9,294 | 237,176 |
| **France** | 4.52 | 161,987 | 49,496 | 74,121 | 285,604 |
| **Gabon** | 9.07 | 11,780 | 118,348 | 120,852 | 250,979 |
| **Gambia** | 4.56 | 181 | 85 | 0 | 266 |
| **Georgia** | 7.79 | 6,982 | 17,803 | 9,784 | 34,570 |
| **Germany** | 2.28 | 122,168 | 11,307 | 0 | 133,475 |
| **Ghana** | 4.53 | 57,519 | 28,901 | 2,160 | 88,580 |
| **Greece** | 6.6 | 14,548 | 27,833 | 1,078 | 43,459 |
| **Grenada** | 4.22 | 221 | 86 | 0 | 308 |
| **Guatemala** | 3.85 | 58,572 | 18,764 | 5,592 | 82,928 |
| **Guinea** | 4.9 | 81,702 | 54,877 | 2,895 | 139,475 |
| **Guinea-Bissau** | 5.7 | 9,274 | 8,702 | 855 | 18,831 |
| **Guyana** | 9.58 | 4,162 | 40,817 | 147,413 | 192,391 |
| **Haiti** | 4.01 | 7,116 | 2,831 | 12 | 9,959 |
| **Honduras** | 4.48 | 57,899 | 23,802 | 3,692 | 85,392 |
| **Hungary** | 2.25 | 18,729 | 2,047 | 0 | 20,776 |
| **India** | 7.09 | 117,992 | 254,792 | 54,428 | 427,211 |
| **Indonesia** | 6.6 | 535,370 | 509,018 | 431,973 | 1,476,361 |
| **Iran** | 7.67 | 3,361 | 12,930 | 2,162 | 18,453 |
| **Iraq** | 3.59 | 104 | 9 | 0 | 113 |
| **Ireland** | 0.92 | 5,283 | 96 | 0 | 5,378 |
| **Israel** | 4.14 | 170 | 85 | 0 | 255 |
| **Italy** | 3.65 | 79,403 | 26,858 | 25 | 106,286 |
| **Jamaica** | 5.01 | 5,362 | 3,249 | 158 | 8,770 |
| **Japan** | 5.8 | 135,783 | 133,480 | 16,005 | 285,268 |
| **Jordan** | 2.79 | 12 | 0 | 0 | 12 |
| **Kazakhstan** | 8.23 | 6,068 | 18,926 | 15,294 | 40,288 |
| **Kenya** | 4.2 | 28,427 | 13,558 | 4,702 | 46,686 |
| **Kosovo** | 5.19 | 2,628 | 1,775 | 47 | 4,450 |
| **Kyrgyzstan** | 8.86 | 329 | 2,819 | 2,761 | 5,909 |
| **Laos** | 5.59 | 92,986 | 80,564 | 19,252 | 192,801 |
| **Latvia** | 2.09 | 38,164 | 2,137 | 0 | 40,301 |
| **Lebanon** | 3.76 | 541 | 115 | 0 | 656 |
| **Lesotho** | 7.4 | 1 | 4 | 0 | 5 |
| **Liberia** | 4.79 | 51,975 | 31,162 | 11,025 | 94,163 |
| **Libya** | 4.85 | 15 | 2 | 0 | 17 |
| **Liechtenstein** | 4.5 | 59 | 42 | 0 | 101 |
| **Lithuania** | 1.62 | 24,554 | 930 | 0 | 25,484 |
| **Luxembourg** | 1.12 | 1,170 | 0 | 0 | 1,170 |
| **Macedonia** | 7.42 | 2,034 | 7,090 | 459 | 9,583 |
| **Madagascar** | 4.63 | 120,340 | 66,584 | 11,922 | 198,846 |
| **Malawi** | 5.74 | 12,514 | 12,167 | 2,396 | 27,078 |
| **Malaysia** | 5.01 | 130,825 | 91,957 | 21,499 | 244,281 |
| **Mali** | 7.16 | 451 | 996 | 140 | 1,586 |
| **Mauritius** | 5.46 | 567 | 478 | 0 | 1,045 |
| **Mexico** | 6.82 | 193,908 | 280,445 | 121,842 | 596,195 |
| **Micronesia** | 7.55 | 8 | 35 | 0 | 43 |
| **Moldova** | 2.2 | 3,113 | 202 | 0 | 3,315 |
| **Mongolia** | 9.36 | 520 | 11,915 | 27,407 | 39,841 |
| **Montenegro** | 6.41 | 2,949 | 4,778 | 82 | 7,809 |
| **Morocco** | 6.74 | 2,260 | 4,076 | 451 | 6,787 |
| **Mozambique** | 6.93 | 150,665 | 189,362 | 115,379 | 455,406 |
| **Myanmar** | 7.18 | 129,745 | 220,188 | 96,924 | 446,857 |
| **Namibia** | 8.43 | 5 | 13 | 17 | 36 |
| **Nepal** | 7.23 | 13,785 | 41,992 | 3,760 | 59,538 |
| **Netherlands** | 0.6 | 5,250 | 72 | 0 | 5,322 |
| **New Zealand** | 7.12 | 34,503 | 44,155 | 35,334 | 113,992 |
| **Nicaragua** | 3.63 | 65,356 | 17,646 | 4,858 | 87,860 |
| **Nigeria** | 6.2 | 64,621 | 65,355 | 24,307 | 154,283 |
| **North Korea** | 8.02 | 8,374 | 40,156 | 8,410 | 56,939 |
| **Norway** | 6.98 | 39,343 | 67,383 | 16,627 | 123,352 |
| **Pakistan** | 7.42 | 2,090 | 7,859 | 1,139 | 11,088 |
| **Palau** | 8.09 | 45 | 333 | 9 | 387 |
| **Panama** | 6.37 | 25,420 | 21,310 | 14,605 | 61,336 |
| **Papua New Guinea** | 8.84 | 37,294 | 183,415 | 216,355 | 437,064 |
| **Paraguay** | 6.39 | 78,538 | 102,626 | 29,877 | 211,041 |
| **Peru** | 8.86 | 85,793 | 190,547 | 509,720 | 786,061 |
| **Philippines** | 5.91 | 91,820 | 100,831 | 8,393 | 201,044 |
| **Poland** | 2.24 | 101,886 | 7,103 | 0 | 108,989 |
| **Portugal** | 8.82 | 25,966 | 553 | 0 | 26,519 |
| **Romania** | 5.95 | 38,395 | 48,394 | 607 | 87,395 |
| **Russian Federation** | 9.02 | 739,484 | 2,245,281 | 5,137,079 | 8,121,843 |
| **Rwanda** | 3.85 | 5,665 | 2,170 | 619 | 8,454 |
| **Saint Kitts and Nevis** | 4.55 | 95 | 50 | 0 | 145 |
| **Saint Lucia** | 6.17 | 235 | 316 | 0 | 551 |
| **Saint Vincent and the Grenadines** | 6.95 | 91 | 221 | 0 | 312 |
| **San Marino** | 0.01 | 7 | 0 | 0 | 7 |
| **Sao Tome and Principe** | 6.64 | 31 | 140 | 0 | 171 |
| **Senegal** | 7.11 | 847 | 2,456 | 162 | 3,465 |
| **Serbia** | 5.29 | 17,513 | 14,112 | 516 | 32,141 |
| **Seychelles** | 10 | 0 | 0 | 68 | 68 |
| **Sierra Leone** | 2.76 | 52,512 | 11,858 | 640 | 65,010 |
| **Singapore** | 1.11 | 170 | 2 | 0 | 172 |
| **Slovakia** | 4.34 | 17,615 | 8,165 | 0 | 25,781 |
| **Slovenia** | 3.78 | 11,065 | 3,791 | 0 | 14,856 |
| **Solomon Islands** | 7.19 | 6,871 | 15,310 | 3,149 | 25,329 |
| **Somalia** | 7.16 | 347 | 1,384 | 46 | 1,777 |
| **South Africa** | 4.94 | 45,489 | 34,968 | 3,196 | 83,653 |
| **South Korea** | 6.02 | 25,060 | 32,009 | 888 | 57,956 |
| **South Sudan** | 9.45 | 5,083 | 59,389 | 146,218 | 210,691 |
| **Spain** | 4.23 | 82,770 | 46,013 | 133 | 128,916 |
| **Sri Lanka** | 5.83 | 20,541 | 22,390 | 1,613 | 44,544 |
| **Sudan** | 9.8 | 1 | 72 | 495 | 569 |
| **Suriname** | 9.39 | 6,796 | 25,031 | 107,954 | 139,781 |
| **Swaziland** | 4.21 | 5,054 | 2,501 | 14 | 7,569 |
| **Sweden** | 5.35 | 174,415 | 109,779 | 23,494 | 307,687 |
| **Switzerland** | 3.53 | 13,636 | 4,412 | 10 | 18,058 |
| **Syria** | 3.64 | 841 | 282 | 0 | 1,123 |
| **Taiwan** | 6.38 | 8,786 | 14,547 | 1,453 | 24,786 |
| **Tajikistan** | 8.65 | 34 | 137 | 130 | 301 |
| **Tanzania** | 7.13 | 123,997 | 159,712 | 122,812 | 406,521 |
| **Thailand** | 6 | 86,276 | 89,326 | 33,612 | 209,214 |
| **Timor-Leste** | 7.11 | 1,783 | 7,008 | 47 | 8,838 |
| **Togo** | 5.88 | 5,064 | 4,522 | 1,076 | 10,662 |
| **Trinidad and Tobago** | 6.62 | 1,478 | 2,176 | 418 | 4,072 |
| **Tunisia** | 5.14 | 1,354 | 987 | 0 | 2,340 |
| **Turkey** | 6.39 | 43,043 | 68,243 | 3,516 | 114,801 |
| **Turkmenistan** | 6.31 | 5 | 33 | 0 | 37 |
| **Uganda** | 4.36 | 77,303 | 36,381 | 7,507 | 121,190 |
| **Ukraine** | 3.3 | 89,540 | 20,183 | 176 | 109,900 |
| **United Kingdom** | 1.65 | 29,149 | 2,917 | 35 | 32,101 |
| **United States** | 6.65 | 1,328,079 | 1,144,693 | 658,645 | 3,131,417 |
| **Uruguay** | 3.61 | 11,793 | 3,998 | 0 | 15,791 |
| **Uzbekistan** | 6.77 | 214 | 227 | 199 | 640 |
| **Vanuatu** | 8.82 | 734 | 5,322 | 4,448 | 10,504 |
| **Venezuela** | 8.78 | 64,650 | 170,792 | 351,112 | 586,554 |
| **Vietnam** | 5.35 | 82,551 | 75,353 | 9,588 | 167,492 |
| **Zambia** | 7.5 | 96,969 | 164,376 | 110,822 | 372,167 |
| **Zimbabwe** | 6.31 | 9,450 | 14,417 | 1,644 | 25,511 |

**Table S6.**
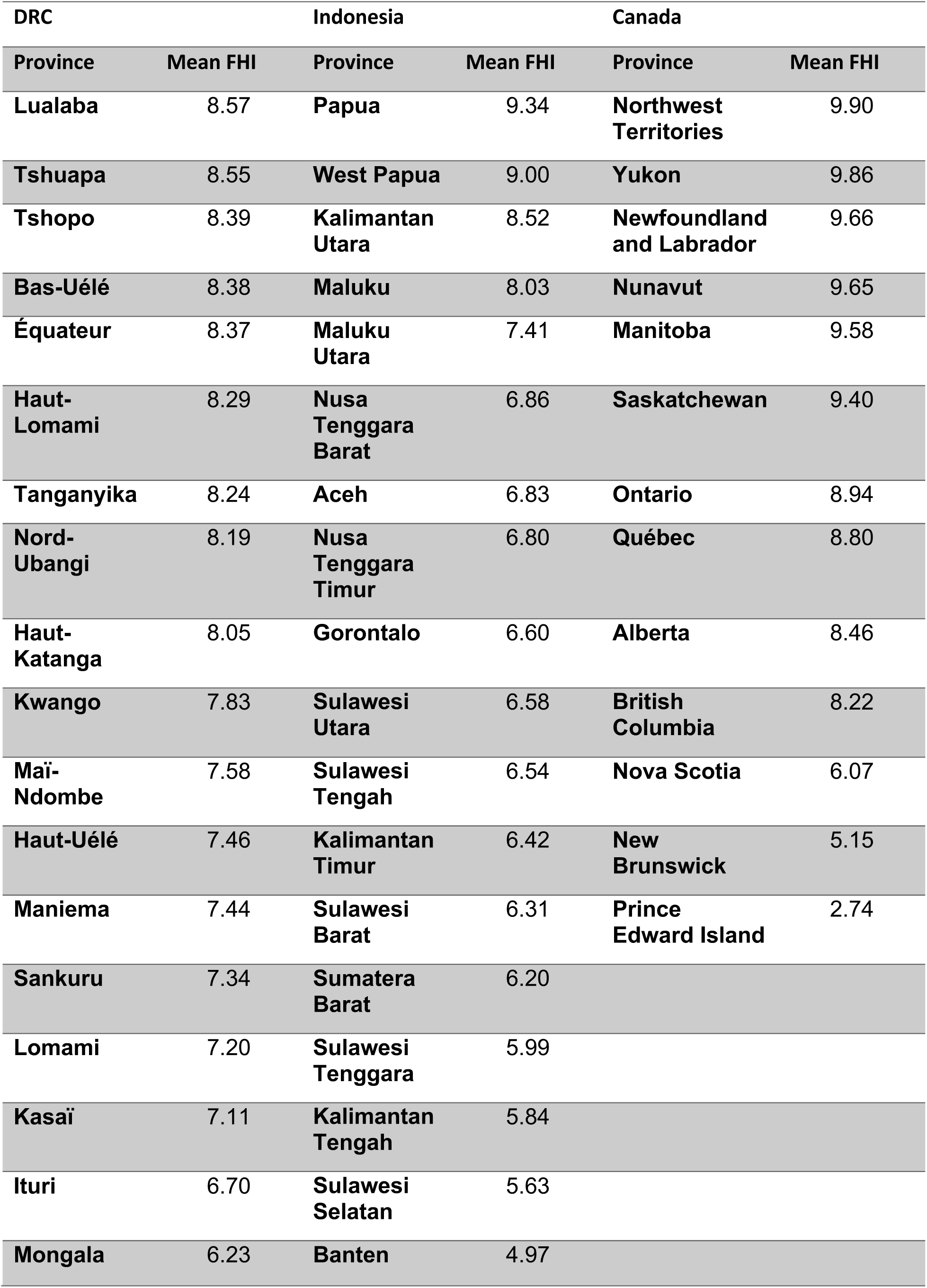
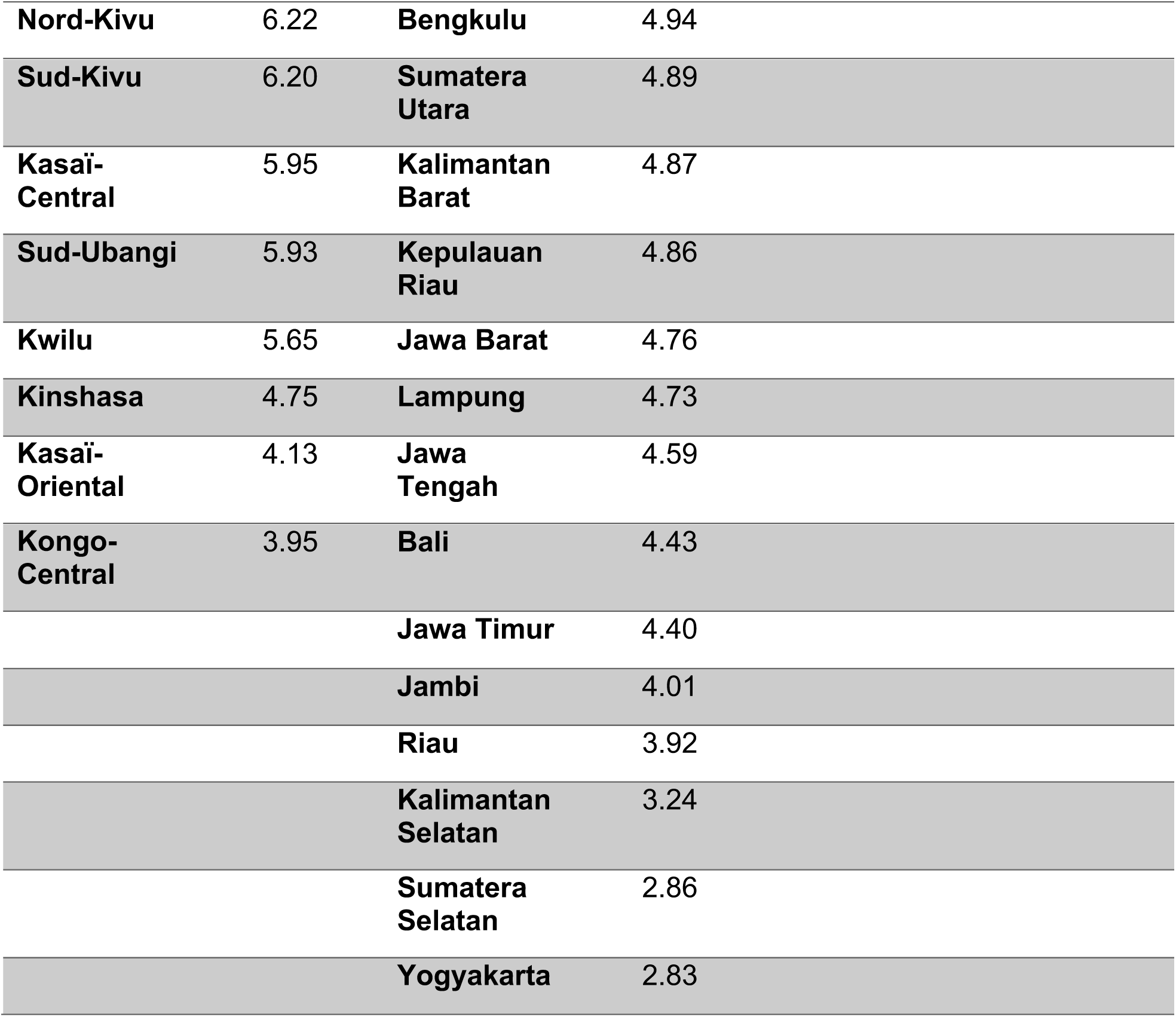
Mean Forest Landscape Integrity Index scores for provinces of Democratic Republic of Congo (DRC), Indonesia and Canada.

| DRC |  | Indonesia |  | Canada |  |
| --- | --- | --- | --- | --- | --- |
| Province | Mean FHI | Province | Mean FHI | Province | Mean FHI |
| Lualaba | 8.57 | Papua | 9.34 | Northwest Territories | 9.90 |
| Tshuapa | 8.55 | West Papua | 9.00 | Yukon | 9.86 |
| Tshopo | 8.39 | Kalimantan Utara | 8.52 | Newfoundland and Labrador | 9.66 |
| Bas-Uélé | 8.38 | Maluku | 8.03 | Nunavut | 9.65 |
| Équateur | 8.37 | Maluku Utara | 7.41 | Manitoba | 9.58 |
| Haut-Lomami | 8.29 | Nusa Tenggara Barat | 6.86 | Saskatchewan | 9.40 |
| Tanganyika | 8.24 | Aceh | 6.83 | Ontario | 8.94 |
| Nord-Ubangi | 8.19 | Nusa Tenggara Timur | 6.80 | Québec | 8.80 |
| Haut-Katanga | 8.05 | Gorontalo | 6.60 | Alberta | 8.46 |
| Kwango | 7.83 | Sulawesi Utara | 6.58 | British Columbia | 8.22 |
| Mai-Ndombe | 7.58 | Sulawesi Tengah | 6.54 | Nova Scotia | 6.07 |
| Haut-Uélé | 7.46 | Kalimantan Timur | 6.42 | New Brunswick | 5.15 |
| Maniema | 7.44 | Sulawesi Barat | 6.31 | Prince Edward Island | 2.74 |
| Sankuru | 7.34 | Sumatera Barat | 6.20 |  |  |
| Lomami | 7.20 | Sulawesi Tenggara | 5.99 |  |  |
| Kasaï | 7.11 | Kalimantan Tengah | 5.84 |  |  |
| Ituri | 6.70 | Sulawesi Selatan | 5.63 |  |  |
| Mongala | 6.23 | Banten | 4.97 |  |  |

|  |  |  |  |
| --- | --- | --- | --- |
| **Nord-Kivu** | 6.22 | **Bengkulu** | 4.94 |
| **Sud-Kivu** | 6.20 | **Sumatera Utara** | 4.89 |
| **Kasaï-Central** | 5.95 | **Kalimantan Barat** | 4.87 |
| **Sud-Ubangi** | 5.93 | **Kepulauan Riau** | 4.86 |
| **Kwilu** | 5.65 | **Jawa Barat** | 4.76 |
| **Kinshasa** | 4.75 | **Lampung** | 4.73 |
| **Kasaï-Oriental** | 4.13 | **Jawa Tengah** | 4.59 |
| **Kongo-Central** | 3.95 | **Bali** | 4.43 |
|  |  | **Jawa Timur** | 4.40 |
|  |  | **Jambi** | 4.01 |
|  |  | **Riau** | 3.92 |
|  |  | **Kalimantan Selatan** | 3.24 |
|  |  | **Sumatera Selatan** | 2.86 |
|  |  | **Yogyakarta** | 2.83 |

**Figure S1.**
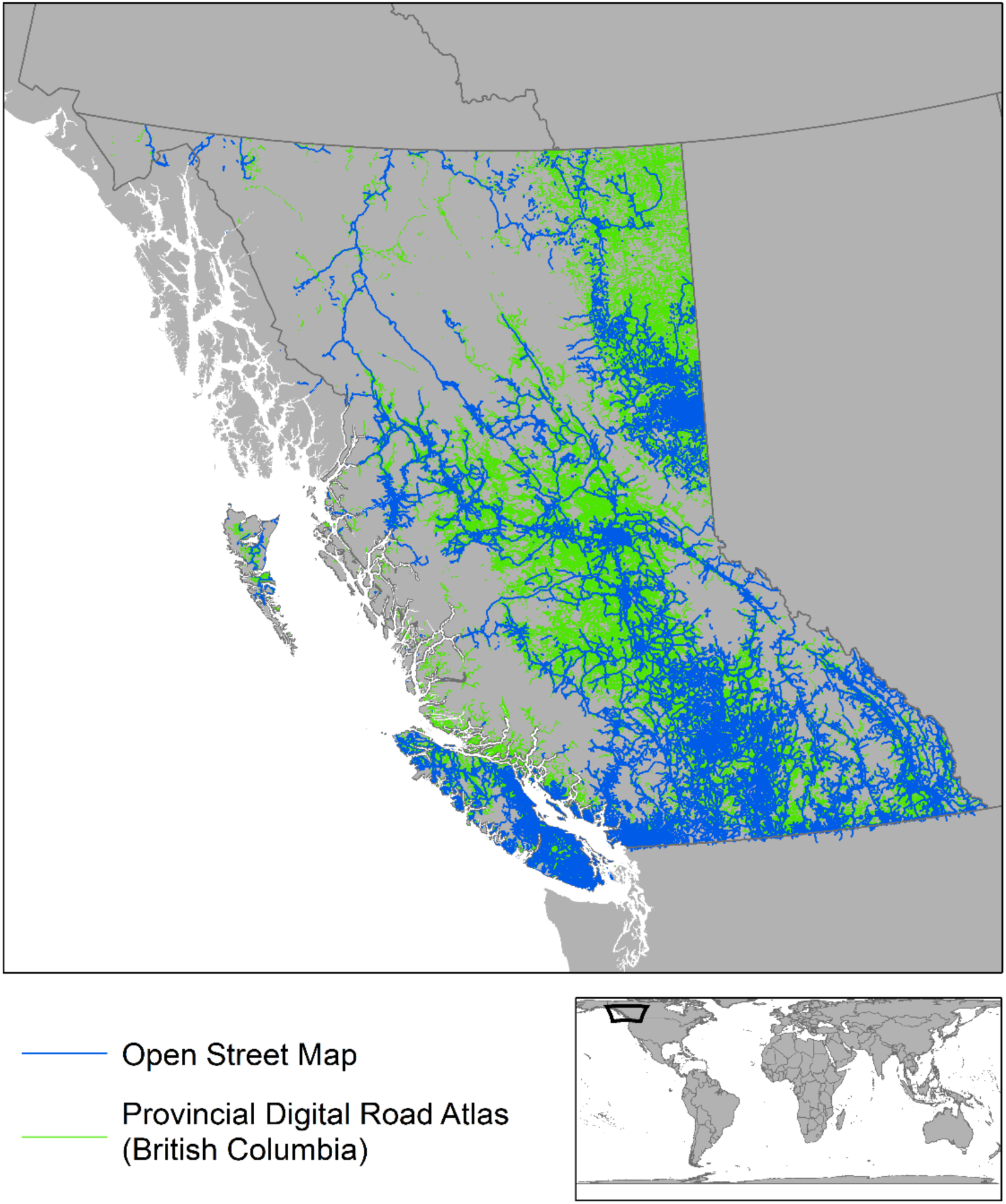
A map overlaying the Open Street Maps data (blue) and provincial government data (green) for roads and other linear infrastructure associated with resource access.

**Figure S2.**
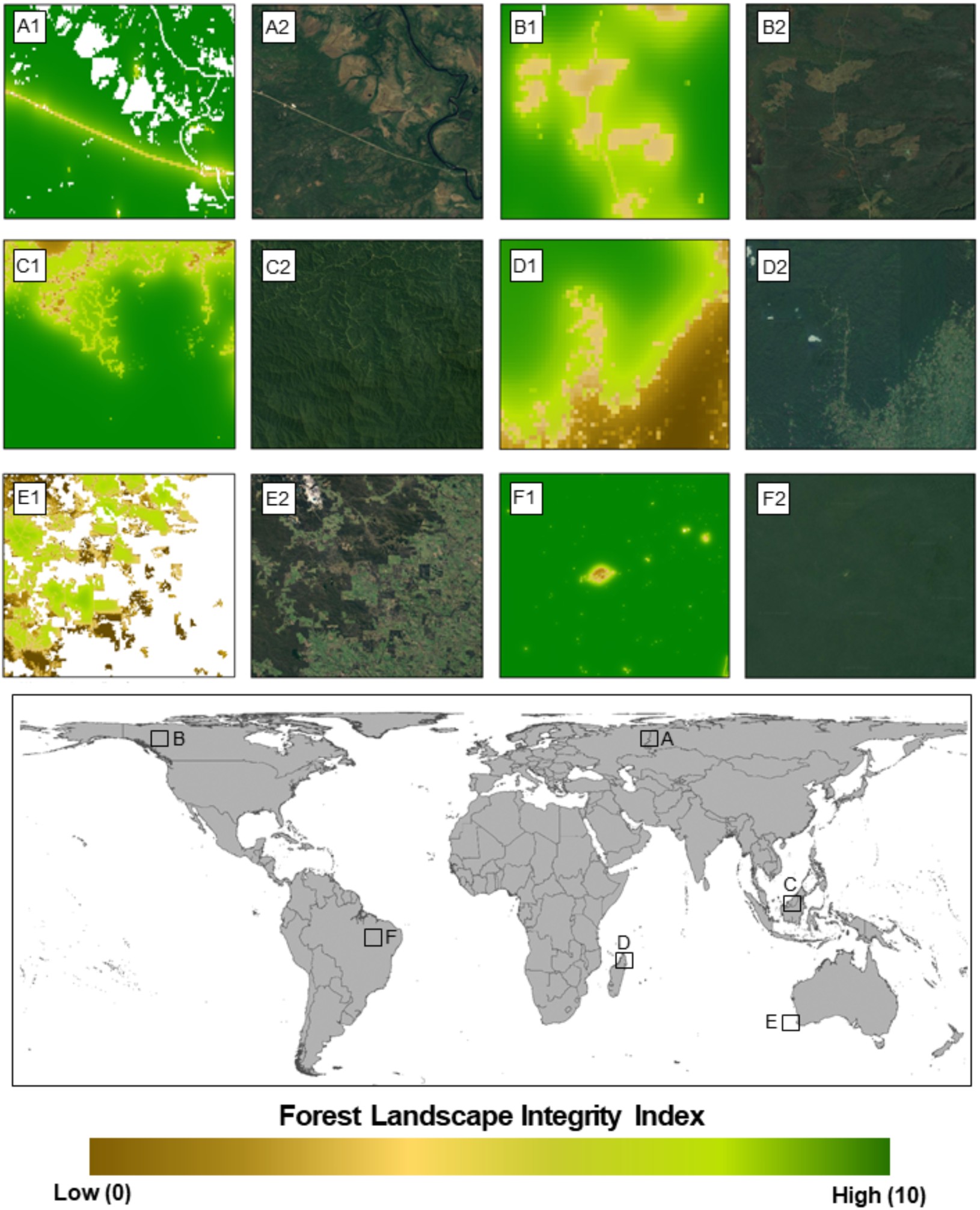
A global map of Forest Landscape Integrity for 2019. Highlighted regions show **A.** A remote road in Russia, **B.** Clearcut logging in Canada, **C.** Selective logging in Borneo, **D.** Swidden agriculture in Madagascar, **E.** Forest fragmentation in Western Australia, **F.** Remote settlements in the Brazilian Amazon.

